## Supplementary Information for "Phenotypic plasticity as a mechanism of cave colonization and adaptation"

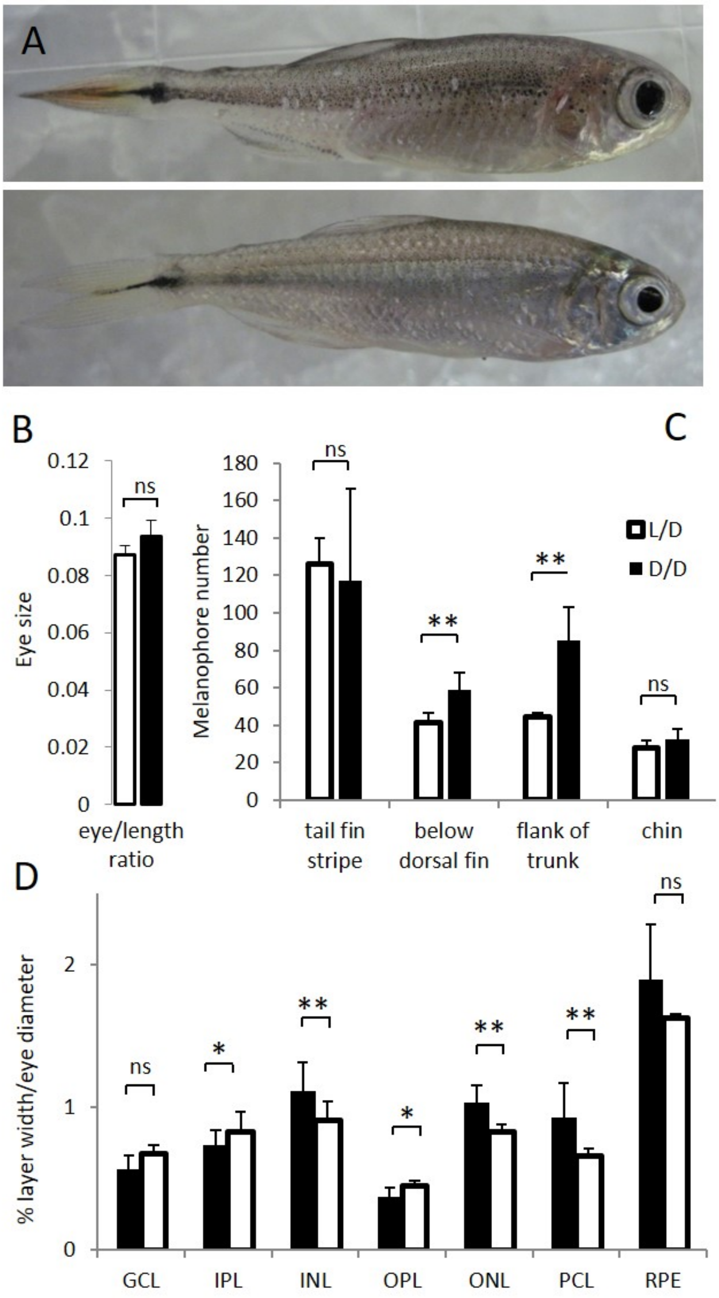


**Figure 1.** A. Surface fish (SF) kept in constant dark (D/D; top frame) vs. light/dark (L/D; bottom frame) photoperiod for 1 year. B. Eye size normalized by body length in D/D vs. L/D SF kept in the experimental conditions for 1 to 2 years. C. Number of melanophores in 1 year-old D/D vs. L/D SF determined in 4 different body regions. D. Thickness of retinal layers in D/D vs. L/D fish: GCL, ganglion cell layer; IPL, inner plexiform layer; INL, inner nuclear layer; OPL, outer plexiform layer; ONL, outer nuclear layer; PCL photoreceptor cell layer; RPE, retinal pigment epithelium measured as a ratio to eye diameter. (Error bars: SD; T-test Ns – not significant, *p<0.05, ** P<0.001).

***Gene Expression Changes in Dark-raised Surface Fish***

To understand the effects of exposure to constant darkness at the molecular level, a comparative transcriptomic analysis was conducted. We used three, 7 month-old surface fish placed in either D/D or L/D conditions within 1 dpf. RNA sequencing yielded 21.9 to 27.6 million reads from each fish, representing a total of 25,194 genes. Basic statistical data are included in Supplementary Table 2. We found 356 differentially expressed genes (DEG) at a significance threshold of $p_{adj}$<0.1 (Supplementary Table 3). Of these, 210 were up-regulated and 146 were down-regulated in the D/D fish; the set of DEGs contained 67 genes with unconfirmed functions. We were particularly interested in the genes related to known aspects of the CF phenotype. Genes involved in circadian regulation, locomotor rhythm and visual perception were down-regulated in D/D fish, whereas lipid metabolism was the main functional category enriched in the up-regulated gene set. Two genes involved in pigmentation were also changed but their expression was higher in D/D, consistent with the increase in pigment cells described above. We also found significant changes in gene expression which were not predicted from known CF-associated phenotypes. These genes function in oxidation-reduction processes, hormone activity, hemostasis, aromatic amino acid metabolism, gene expression, metabolism of proteins, and signal transduction (Fig 2).

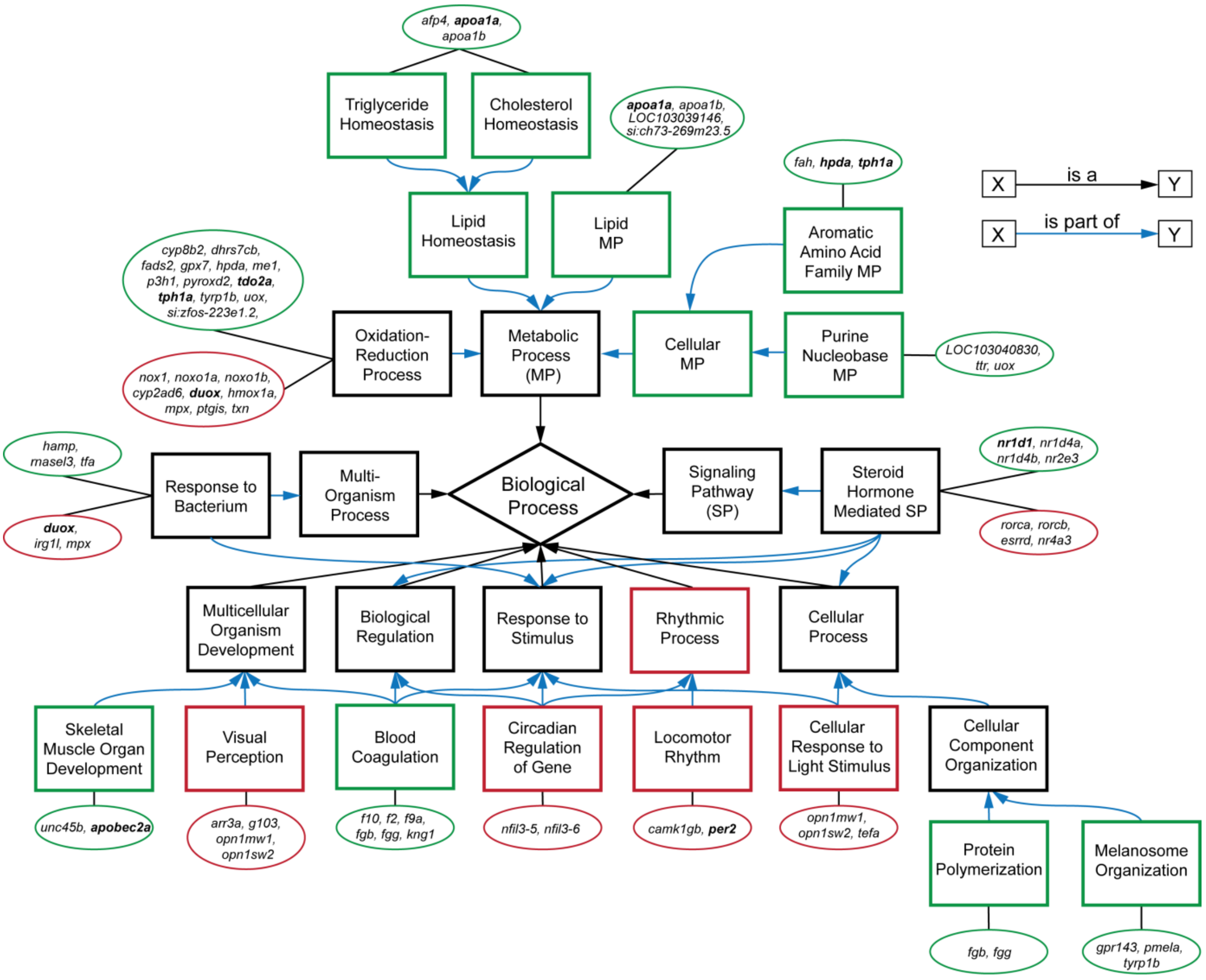

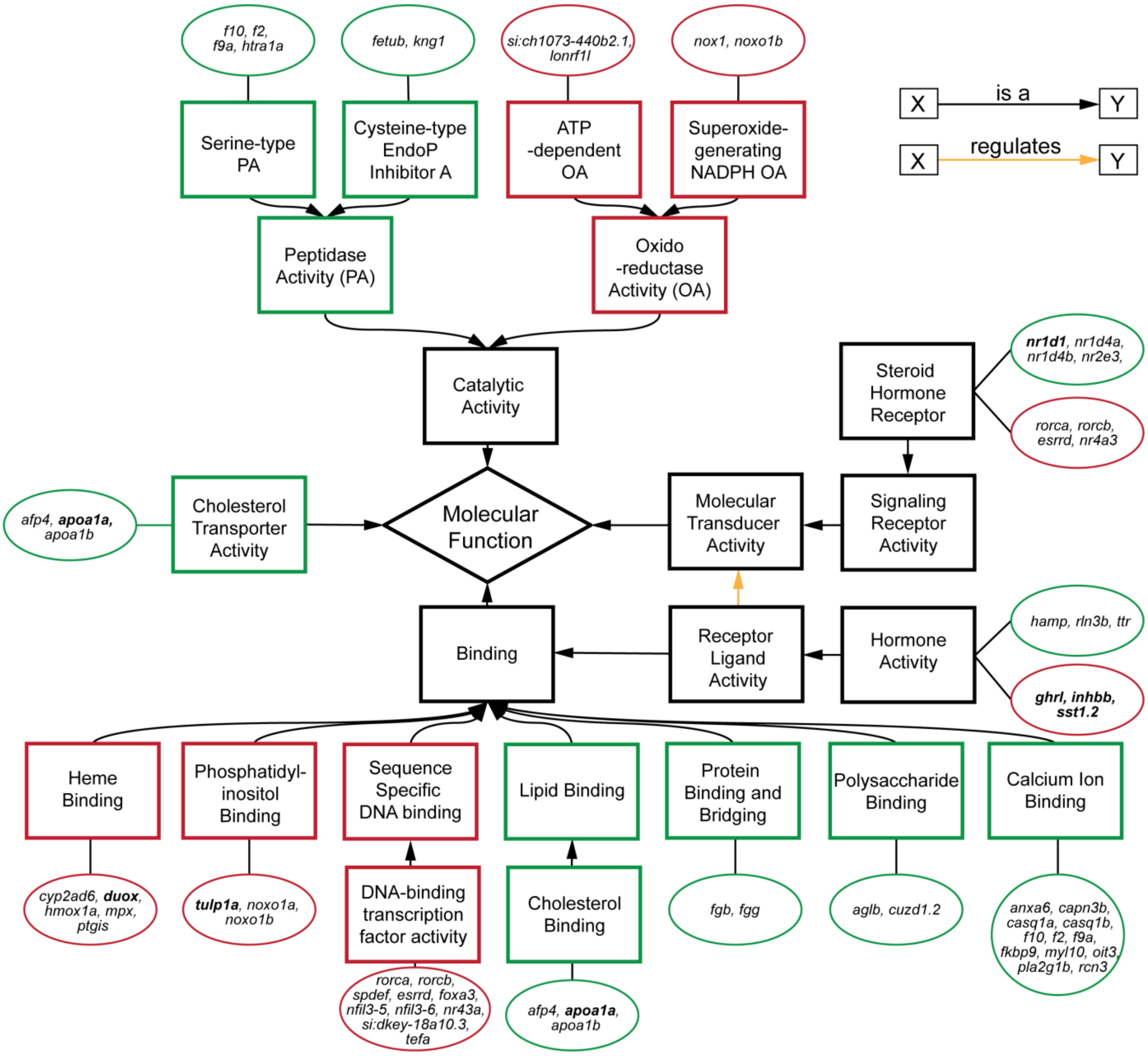


**Figure 2**. Subset of relevant, enriched GO terms (boxes) for biological processes (top) and molecular functions (bottom) of DEGs (circles) in the transcriptome. Genes tested by RT-PCR are in bold. Red outlines are down-regulated terms and genes, and green outlines are up-regulated.

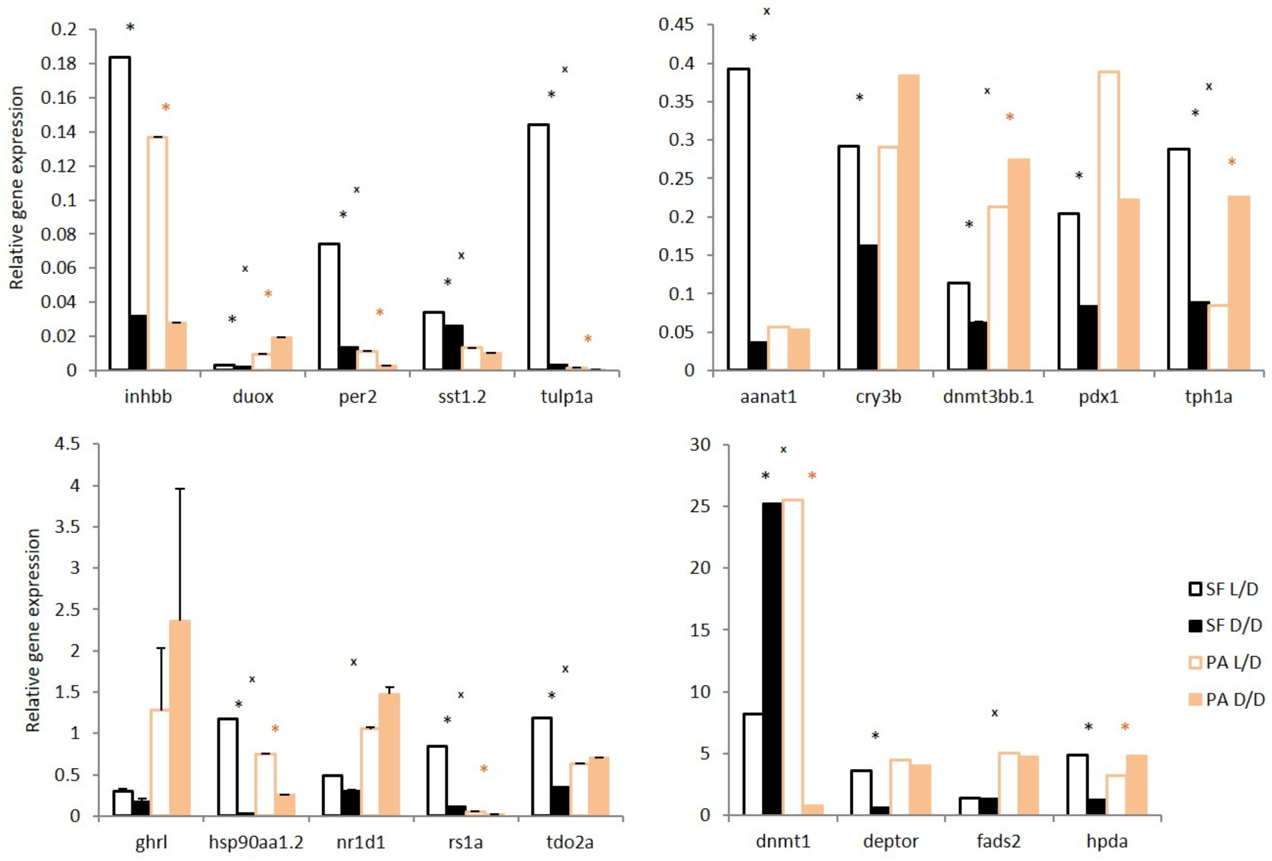


**Figure 3**. Normalized relative expression levels of genes in D/D and L/D SF and PA determined by RT-PCR. (Error bars: SD; ANOVA with Bonferroni adjustments $p<0.05$ * for SF D/D vs. SF L/D; * for PA D/D vs. PA L/D; $^{X}$ for SF vs. PA,).

### *Starvation Resistance in Dark-raised Surface Fish*

A major challenge facing animals that colonize caves is low food availability due to limited or absent primary productivity. To determine how a surface ancestor may have coped with this difficulty, we raised SF and PA larvae in D/D and L/D beginning <24 hours post fertilization (hpf) without feeding (N=36 larvae/group). D/D and L/D controls were fed daily portions of brine shrimp from 7 dpf, when larvae normally lose their reliance on yolk and begin feeding. By 15 dpf, about 25% more SF and over 65% more PA unfed larvae were alive in D/D compared to L/D conditions. By 18 dpf, only a few unfed embryos were still alive, and they were all from the D/D conditions (8 PA and 2 SF) (Fig 4). The results show that dark raised SF and CF are more resistant to starvation than siblings raised under a normal photoperiod and that PA larvae survive starvation better than SF larvae regardless of the lighting conditions.

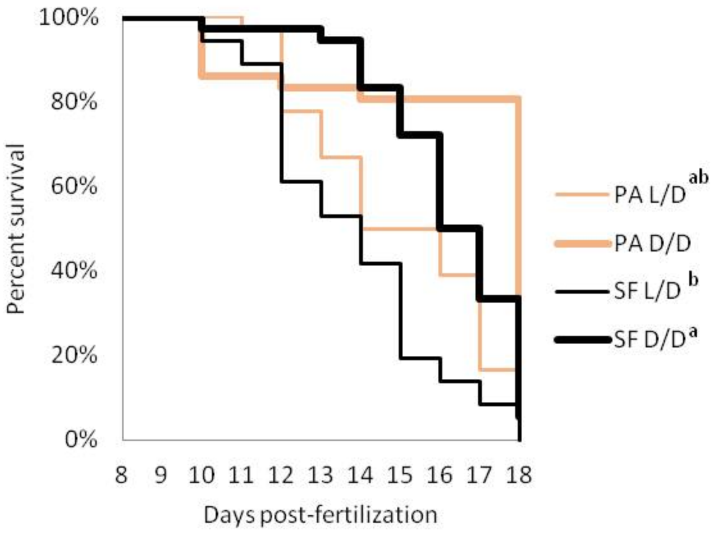


**Figure 4.** Survival curve of starvation resistance in *Astyanax mexicanus* surface fish (SF) and Pachón cavefish (PA) raised in complete darkness (D/D) or a normal photoperiod (L/D). Graphs show the percent of surviving fish (from the initial 36) on each day. Groups of SF and PA larvae from each condition were starved starting at 7 dpf. Vertical drops represent individuals lost at a given time point. Groups in the legend that share a superscript are not statistically different, p values calculated by Cox proportional hazards model followed by generalized linear hypothesis test.

### *Metabolic Rate Decrease in Dark-raised Surface Fish*

We hypothesized that fish in D/D survived starvation longer because of lower energy expenditure. To test this possibility, we exposed SF and PA to D/D vs. L/D conditions within first 24 hpf after spawning and measured $O_2$ consumption at 2.5 dpf and 7.5 dpf. The results showed that SF and PA larvae raised in D/D have decreased metabolic rates when compared to SF and PA larvae raised in L/D conditions. This result was found to be significant in SF each time this experiment was conducted (3 replicates) for both 2.5 and 7.5-day-old larvae. Furthermore, D/D PA showed a decrease in $O_2$ consumption at 2.5 but not at 7.5 dpf when compared to larvae raised in L/D. In addition, PA and SF metabolic rates were not significantly different at 2.5 dpf, whereas at 7.5 dpf PA has a lower metabolic rate than SF larvae raised in L/D conditions. At 2.5 dpf metabolic rate is similar between SF and PA, and darkness caused it to be reduced in both types of fish. However, by 7.5 dpf the metabolic rate in PA was reduced compared to L/D SF, and PA did not show a plastic response to darkness, whereas at the same age SF metabolic rate was still affected by darkness (Fig 5). In summary, the results support the hypothesis that D/D fish survive starvation longer because of lower metabolic rates.

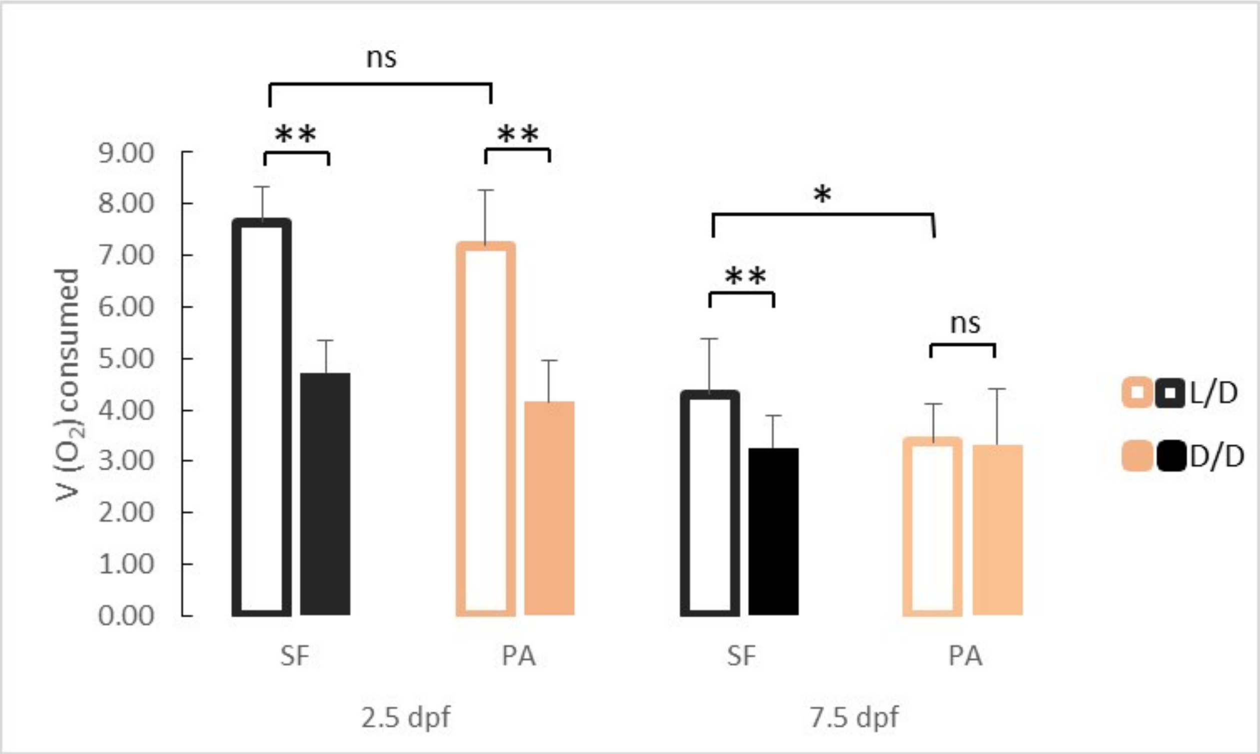


**Figure 5.** Average oxygen consumption at 2.5 and 7.5 dpf in SF and PA larvae kept in D/D versus L/D conditions (N = 18 - 25 larvae/group). (Error bars represent standard deviation, ns: not significant, *p<0.05, **p<0.01 as calculated by ANOVA and Tukey HSD Test.)

### *Cortisol Increases in Dark-raised Surface Fish*

We hypothesized that exposure to constant darkness may represent a chronic stressor and tested this possibility by comparing cortisol levels in both L/D and D/D SF and PA. Surface fish raised in D/D conditions had significantly higher cortisol levels than L/D SF (Fig 6). In contrast, PA did not show a significant change in cortisol levels after exposure to D/D. This result was confirmed in three biological replicates using fish of different ages and different periods of dark exposure and shows that dark raised SF exhibit higher cortisol levels than SF raised on a normal photoperiod.

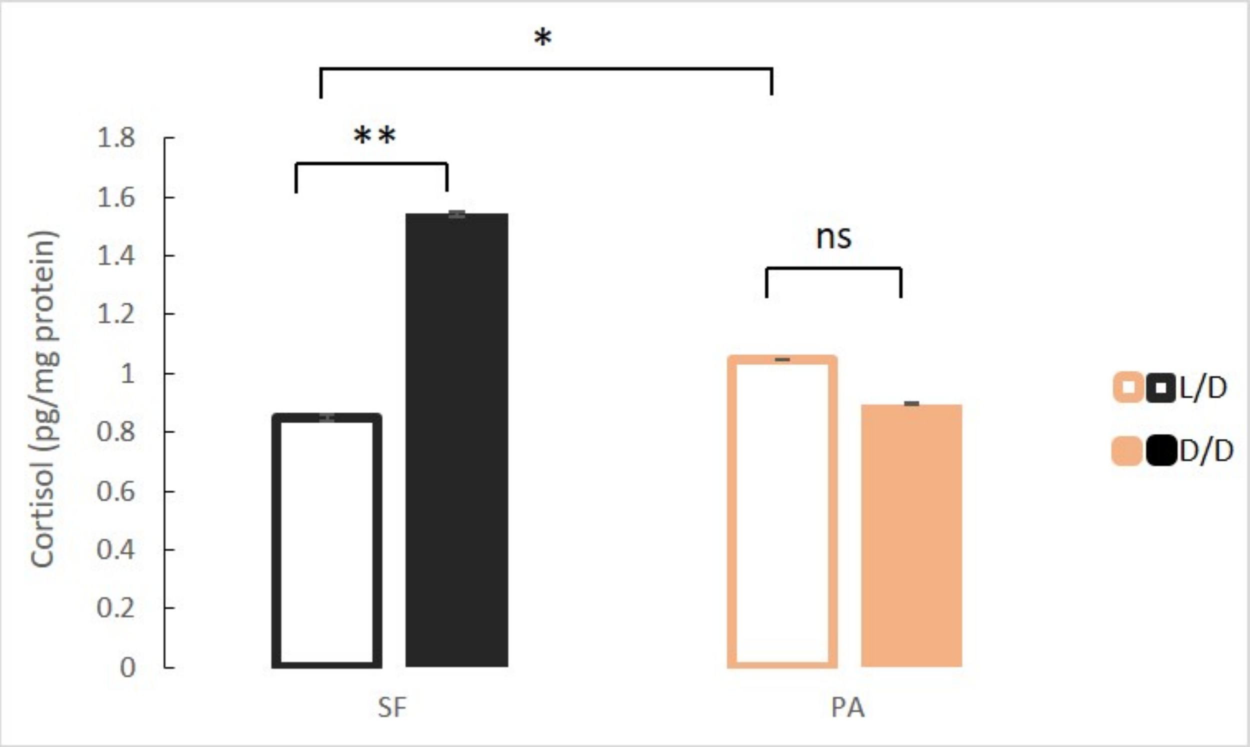


**Figure 6.** Mean cortisol levels in adult surface fish (SF) and Pachón cavefish (PA) kept in D/D or L/D conditions for 1.5 to 2 years (N=4/group). (Error bars represent SD in three technical replicates, ANOVA and Tukey HSD Test: Ns – not significant, *p<0.05, **p<0.01).

***Fat Increase in Dark-raised Surface Fish***

Our RNAseq results show that genes involved in many aspects of fat metabolism were up-regulated in D/D versus L/D reared SF. To confirm this on a phenotypic level, we quantified triglyceride content in SF and PA raised to adulthood under D/D and L/D conditions. In fish raised under L/D conditions, PA had higher triglyceride levels than SF.  Surface fish raised in D/D conditions had markedly higher triglyceride levels than L/D SF. Triglyceride levels in D/D PA were also higher than levels measured in L/D PA, although the difference is more modest than that observed for SF (Fig 7). These results suggest that dark raised SF have higher levels of triglyceride metabolism that SF raised on a normal photoperiod.

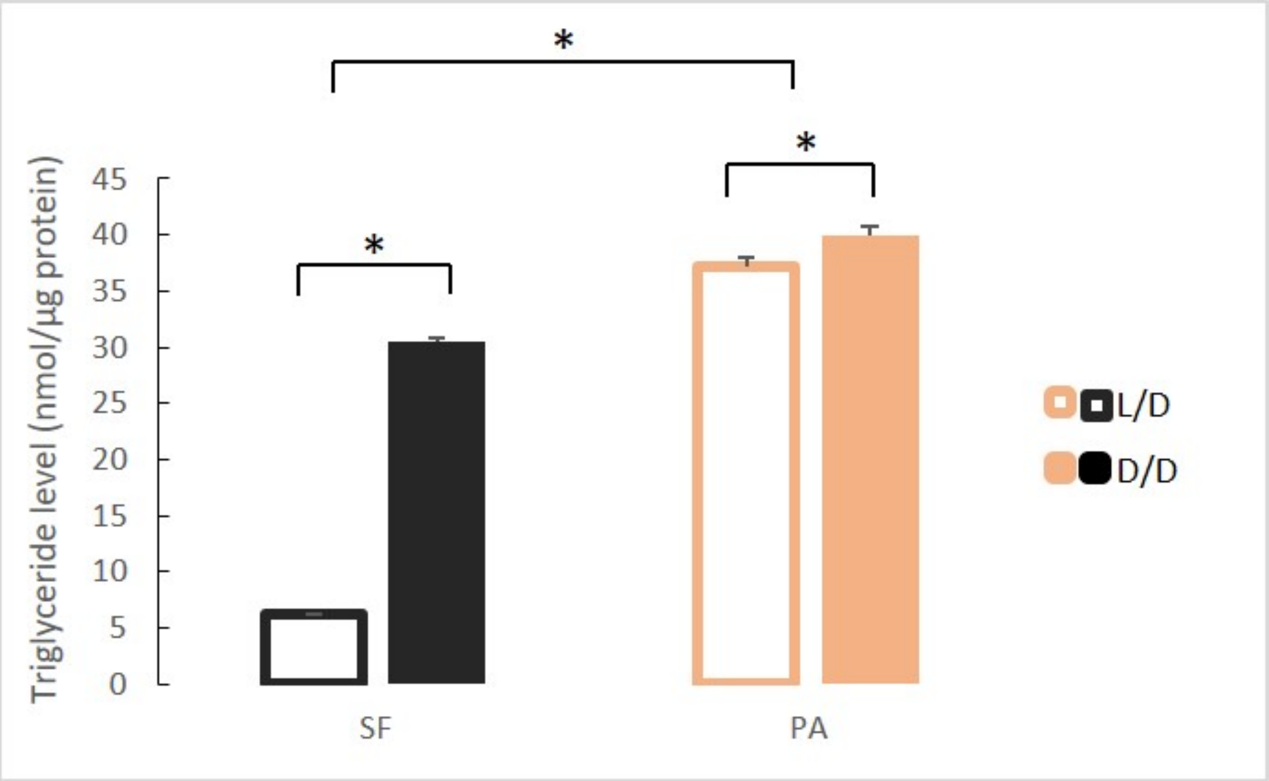


**Figure 7**. Mean triglyceride levels in SF and PA raised under D/D versus L/D conditions for approximately 1 year since < 24 hpf. (Error bars represent standard deviation. *p<0.01; ANOVA and Tukey HSD Test).

***Hormone Levels Change in Dark-raised Surface Fish***

Because of morphological changes in the pituitary and thyroid glands in dark raised SF (47) and differential expression of some of the genes associated with pituitary hormones (e.g. *sst1.2*, *inhbb*, *ghrl*) in the dark-raised SF transcriptome, we quantified the levels of pituitary hormones TSH and GH in SF and PA adults raised in either L/D or D/D conditions. TSH levels were lower in D/D SF (p<0.05) when compared to L/D SF. In contrast, TSH levels in PA raised in L/D and D/D conditions were similar, and PA had higher TSH levels than SF regardless of lighting conditions (Fig 8A). When TSH levels were compared between SF and three populations of CF, there was a trend for higher TSH levels in TI (P=0.07), but not MO (P=0.16), and TSH levels where significantly higher in PA (Fig 8B). The levels of GH were higher in both SF and PA raised in D/D conditions when compared to fish raised in L/D (Fig 9A). All three populations of CF had higher GH levels than SF (Fig 9B).

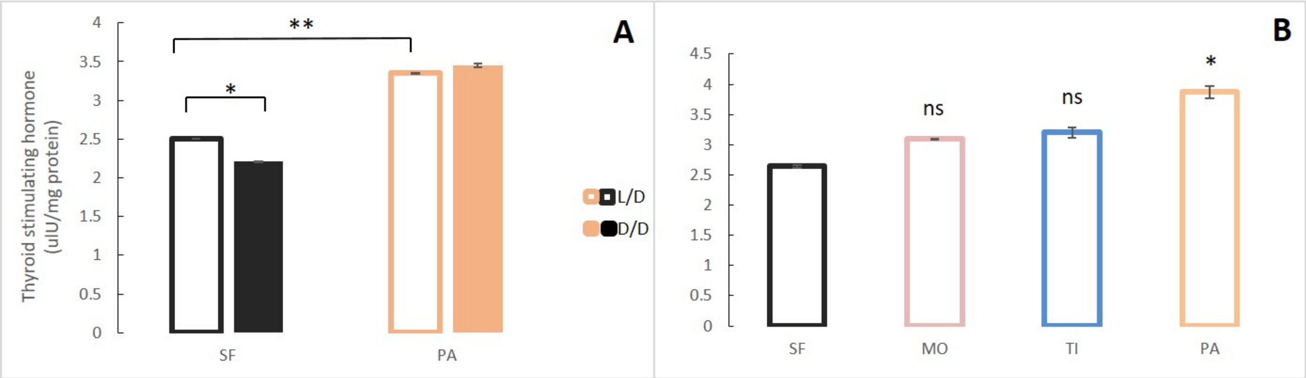


**Figure 8.** A. Mean Thyroid stimulating hormone (TSH) levels normalized by protein concentration in adult surface fish (SF) and Pachón cavefish (PA) kept in D/D or L/D conditions for 1.5 to 2 years. B. Mean TSH levels in SF and 3 different CF populations: Pachón (PA), Tinaja (TI) and Molino (MO) caves. (Error bars represent SD in three technical replicates. N ranges from 3-8 fish/group; * p<0.05; ** p<0.01 as calculated by ANOVA and Tukey HSD Test. In B. ns or * denotes significance in comparison to SF.)

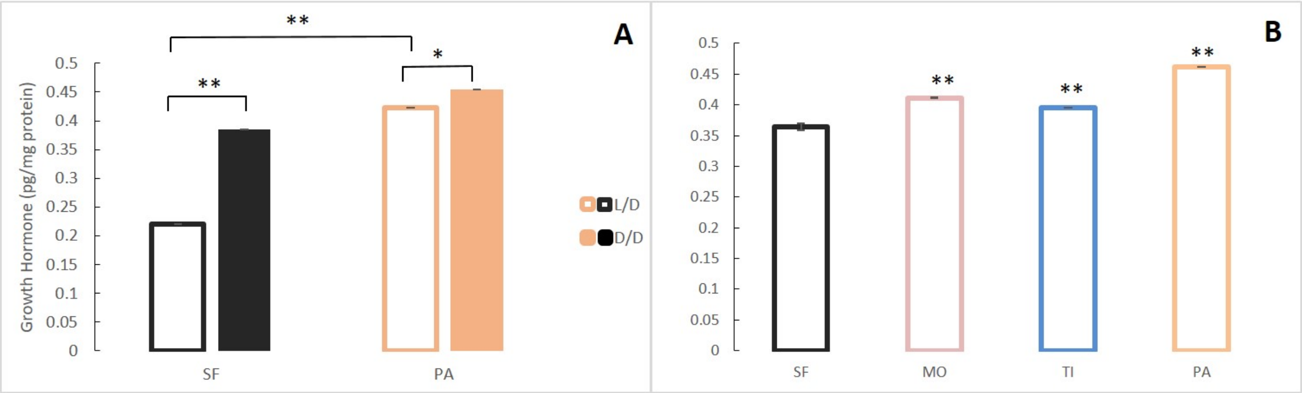


**Figure 9.** A. Mean growth hormone (GH) levels normalized by protein concentration in adult surface fish (SF) and Pachón (PA) cavefish kept in D/D or L/D conditions for 1.5 (SF) and 2 years (PA) since < 3 dpf. B. Mean GH levels in 3-4 month old SF and 3 different CF populations: PA, Tinaja (TI), and Molino (MO). (Error bars represent SD in three technical replicates. N = 3 to 8/group. *p<0.05; **p<0.01 as calculated by ANOVA and Tukey HSD Test. In B, **p<0.01 compared to SF).

### *Serotonin Decrease in Dark-raised Surface Fish*

According to the RNAseq analysis, the tryptophan metabolism pathway was significantly up-regulated in D/D versus L/D SF, while real time qPCR suggested lower expression of the *tdo2a*, *aanat1* and *tph1a* genes in dark–raised SF. Therefore, we compared serotonin (5-HT) levels in D/D and L/D SF and PA adults by HPLC. Since serotonin levels fluctuate according to a daily rhythm (48), samples for this assay were collected between 11 AM and 3 PM (day) or 11 PM and 3 AM (night) regardless of rearing conditions. Serotonin concentrations were lower in the brains (Fig 10A) and the bodies (Fig 10B) of D/D compared to L/D SF. Likewise, the 5-HT metabolite 5-HIAA was lower in the brain of D/D SF (Supplementary Fig 2). It was not possible to quantify 5-HIAA in the body because it was masked by several interfering peaks. Serotonin levels in the SF brain, but not the body, were significantly lower at night when compared to samples collected during the day. In PA brains collected during the day, 5-HT levels were lower in fish raised in D/D when compared to L/D conditions but 5-HT levels were similar in D/D and L/D brains at night. Modest, but significant, changes in body 5-HT were evident in PA with increases in D/D relative to L/D collected during the day, and the opposite change occurred in bodies collected during the night. Serotonin levels were significantly lower in PA brains and bodies when compared to SF regardless of experimental conditions (D/D, L/D, day/night). Comparison of SF brain 5-HT with TI, MO and PA fish showed that all three cave populations have lower brain 5-HT (Fig 10C). In contrast to adults, 5-HT was significantly higher in PA compared to SF at 7 dpf, and darkness did not affect 5-HT levels in either larval fish types (Fig 10D).

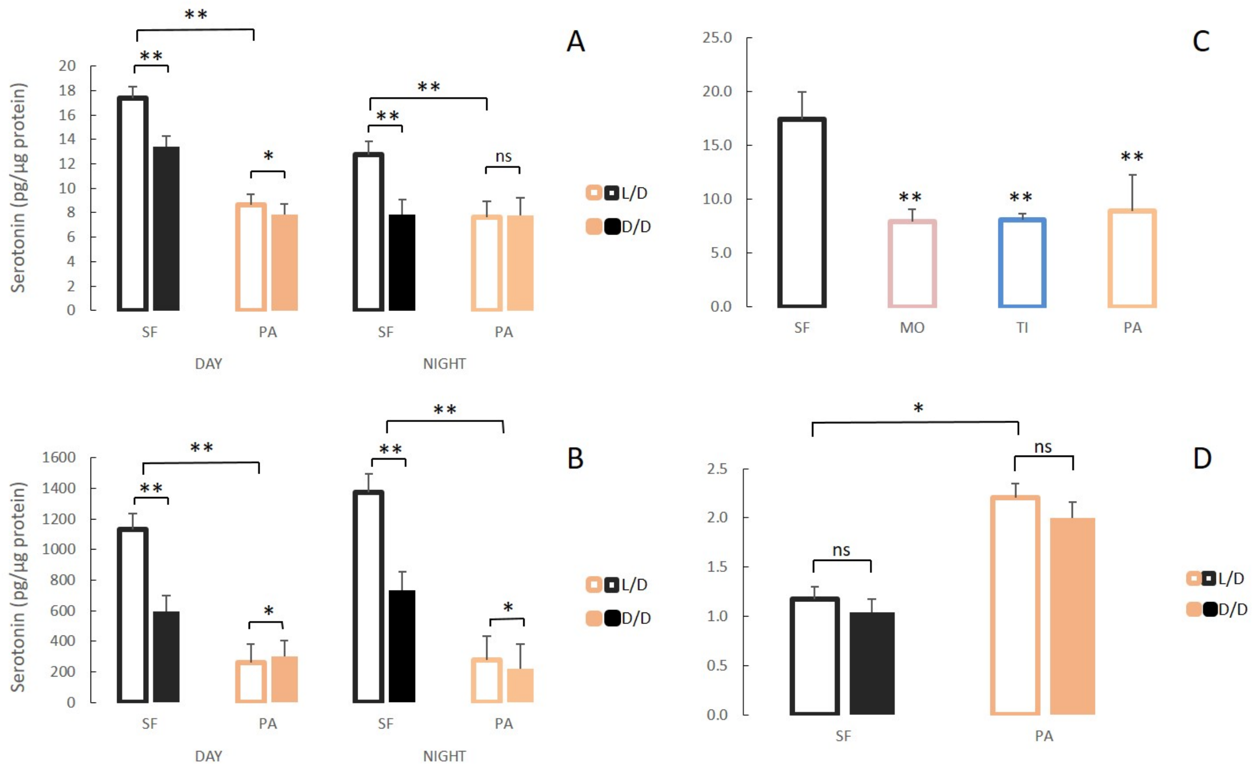


**Figure 10.** Serotonergic system changes in adults and larvae of light/dark (L/D)- and dark/dark (D/D)-reared surface fish and cavefish. A, B. Serotonin levels in adult brains (A) and bodies (B) of D/D and L/D reared surface fish (SF) and Pachón cavefish (PA) collected in the middle of the day (DAY) and the middle of the night (NIGHT). (Error bars represent the standard error of the means.) C. Mean serotonin levels in brains of adult SF and 3 different CF populations: Molino (MO), Tinaja (TI) and PA. D. Mean serotonin levels in pooled samples of 5 larvae aged 7 dpf placed in the experiment within first few hours post fertilization. (Error bars SEM; ns – not significant, *p<0.05; **p<0.01 as calculated by ANOVA and post-

hoc Tukey HSD Test. In C, **p<0.01 vs SF. The number of each fish type subjected to analysis ranged from 4 to 10 per group.)

### *Progeny of Dark Raised Surface Fish Show Plasticity in Starvation Resistance and Metabolic Rate*

To test whether the plastic changes in the fish reared in the dark during the first generation are maintained and transferred to the next generation, we induced spawning in D/D reared SF. We measured starvation resistance and metabolic rate in F1 embryos (dSF) developing in D/D or in L/D during the second generation, as well as the offspring of L/D reared controls exposed to L/D or D/D conditions. The F1 larvae reared in L/D showed no differences in starvation resistance (Fig 11) or metabolic rate (Fig 12), regardless of whether they were the progeny of L/D or D/D parents. Their cohorts raised in darkness showed a modification in these phenotypes: dSF larvae derived from D/D parents showed slightly enhanced plasticity and a minor shift toward lower metabolic rate and higher starvation resistance (although not statistically significant) than larvae raised from L/D parents.

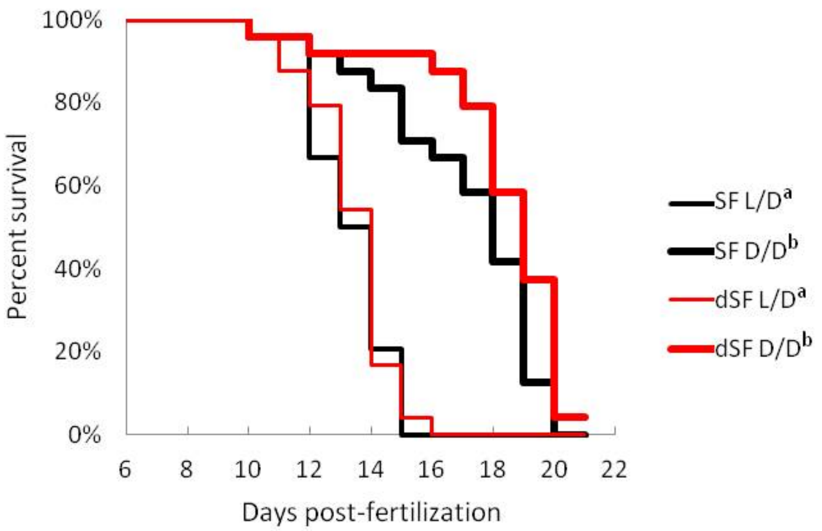


**Figure 11.** Survival curve of starvation resistance in F1 offspring of surface fish kept in normal light/dark photoperiod (SF) and surface fish raised in total darkness for 2 years (dSF). Graphs show the percent of surviving fish (from the initial 24) on each day. A. One group of larvae from each fish type (SF, dSF) and each lighting condition (D/D, L/D) was starved starting at 7 dpf (a vs. b p<0.0001). Vertical drops represent individuals lost at a given time point, groups in the legend that share a superscript are not statistically different, p values calculated by Cox proportional hazards model followed by generalized linear hypothesis test.

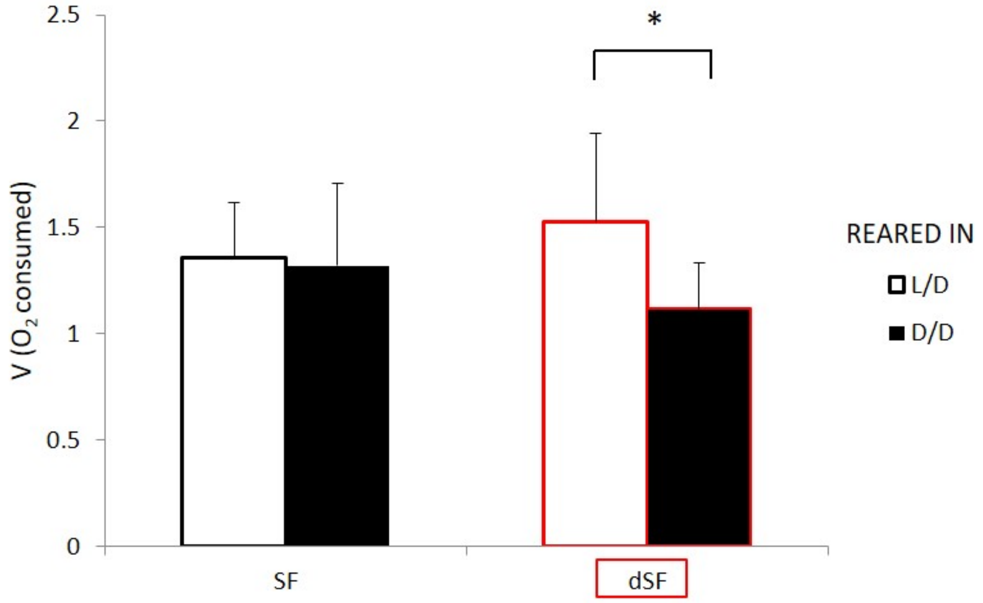


**Figure 12.** Average oxygen consumption of 11 dpf F1 offspring surface fish kept in the normal light/dark photoperiod (SF) and surface fish kept in total darkness for 2 years (dSF). Each group of offspring was exposed to D/D or L/D conditions within first 24 hpf. (Error bars represent standard deviation; *$p<0.05$, as calculated by ANOVA and Tukey HSD Test.).

8. Bradic M, Beerli P, García-de León FJ, Esquivel-Bobadilla S, Borowsky RL. Gene flow

and population structure in the Mexican blind cavefish complex (*Astyanax mexicanus*). BMC Evol Biol [Internet]. 2012 [cited 2013 Jul 11];12(1):9. Available from: http://www.biomedcentral.com/1471-2148/12/9/

19. Hinaux H, Poulain J, Da Silva C, Noirot C, Jeffery WR, Casane D, et al. De Novo

Sequencing of Astyanax mexicanus Surface Fish and Pachón Cavefish Transcriptomes Reveals Enrichment of Mutations in Cavefish Putative Eye Genes. PLoS One [Internet]. 2013 Jan [cited 2013 Jun 14];8(1):e53553. Available from: http://dx.doi.org/10.1371/journal.pone.0053553

http://genomebiology.biomedcentral.com/articles/10.1186/gb-2013-14-4-r36

31. Trapnell C, Williams BA, Pertea G, Mortazavi A, Kwan G, van Baren MJ, et al. Transcript assembly and quantification by RNA-Seq reveals unannotated transcripts and isoform switching during cell differentiation. Nat Biotechnol [Internet]. 2010 May 2 [cited 2019 Feb 15];28(5):511–5. Available from: http://www.nature.com/articles/nbt.1621

32. Trapnell C, Roberts A, Goff L, Pertea G, Kim D, Kelley DR, et al. Differential gene and transcript expression analysis of RNA-seq experiments with TopHat and Cufflinks. Nat Protoc [Internet]. 2012 Mar 1 [cited 2019 Feb 15];7(3):562–78. Available from: http://www.nature.com/doifinder/10.1038/nprot.2012.016

33. Goff LA, Trapnell C, Kelley D. CummeRbund: Visualization and exploration of Cufflinks high-throughput sequencing data [Internet]. 2014 [cited 2019 Feb 15]. Available from: https://www.bioconductor.org/packages/devel/bioc/vignettes/cummeRbund/inst/doc/cummeRbund-manual.pdf

34. R Studio T. RStudio: Integrated development environment for R [Internet]. Citeseer. Boston, MA; 2015 [cited 2019 Feb 15]. Available from: http://www.rstudio.com/.

35. Team R. R: A language and environment for statistical computing. 2018 [cited 2019 Feb 15]; Available from: ftp://ftp.uvigo.es/CRAN/web/packages/dplR/vignettes/intro-dplR.pdf

36. Huang DW, Sherman BT, Lempicki RA. Systematic and integrative analysis of large gene lists using DAVID bioinformatics resources. Nat Protoc [Internet]. 2009 Jan 1 [cited 2019 Feb 15];4(1):44–57. Available from: http://www.nature.com/articles/nprot.2008.211

37. Huang DW, Sherman BT, Lempicki RA. Bioinformatics enrichment tools: paths toward the comprehensive functional analysis of large gene lists. Nucleic Acids Res [Internet]. 2009 Jan 1 [cited 2019 Feb 15];37(1):1–13. Available from: https://academic.oup.com/nar/article-lookup/doi/10.1093/nar/gkn923

38. Fabregat A, Sidiropoulos K, Garapati P, Gillespie M, Hausmann K, Haw R, et al. The Reactome pathway Knowledgebase. Nucleic Acids Res [Internet]. 2016 Jan 4 [cited 2019 Feb 15];44(D1):D481–7. Available from: https://academic.oup.com/nar/article-lookup/doi/10.1093/nar/gkv1351

39. Xie F, Sun G, Stiller JW, Zhang B. Genome-wide functional analysis of the cotton transcriptome by creating an integrated EST database. PLoS One [Internet]. 2011 [cited 2019 Feb 15];6(11):e26980. Available from: http://www.ncbi.nlm.nih.gov/pubmed/22087239

40. Therneau TM, Grambsch PM. Modeling survival data: Extending the Cox model. New York, NY: Springer New York; 2000. (Statistics for Biology and Health).

62. Aspiras AC, Rohner N, Martineau B, Borowsky RL, Tabin CJ. Melanocortin 4 receptor

mutations contribute to the adaptation of cavefish to nutrient-poor conditions. Proc Natl Acad Sci U S A [Internet]. 2015 Aug 4 [cited 2019 Feb 26];112(31):9668–73. Available from: http://www.ncbi.nlm.nih.gov/pubmed/26170297

73. Tarboush R, Chapman GB, Connaughton VP. Retinal development and photoreceptor

synaptic ultrastructure are altered by abnormal rearing light regimes [Internet]. Vol. 20, Eur. J. Anat. 2016 [cited 2019 Jul 24]. Available from: https://pdfs.semanticscholar.org/4b67/f2f0acc031725f5a1a78025a33fd8e2c5c40.pdf

Supplementary Table 1. List of genes and primers used in RT-PCR experiments.

| Gene | Ensembl accession # | Primer A | Primer B |
|---|---|---|---|
| hsp90aa1.2 | ENSAMXG00000020572 | GCCAGACAATGGTGAGTCTATC | GATGCGCTTCTCCTCTGTATATT |
| tdo2a | ENSAMXG00000032882 | CCACCGGATCGTGATGATATT | AGGAGAGAGATACTCCCTGAAG |
| tph1a | ENSAMXG00000018104 | CTTGCCACTTGTTACTTCTTCAC | CAGTTCACTGATGGAGGACAG |
| aanat1 | ENSAMXG00000015728 | CCTTCATCATTGGCTCCCTAT | CACATCAGGATGGAGCCTTT |
| sst1.2 (1 of many) | ENSAMXG00000006540 | TGCACAGGAATGGAAGAAGAG | CCACTCGGACATCCTGTTTAG |
| inhbb | ENSAMXG00000003122 | CGGAGACAGATGACTCAACATT | CCAGGCAACAGCTTGAAGTA |
| rs1a | ENSAMXG00000033915 | CAGGAGGGAGTAGAGGACAAT | CCAACTAGAACTAGCGGTGATG |
| hpda | ENSAMXG00000015913 | CTGAAGACCCTGCAGGAATTA | CTCTGGATGACCTCTAGGAAGA |
| cry3b | ENSAMXG00000008895 | ATTTCGAGAGGCCCAAGATG | TGCTTCACCCGTTTGTAGAG |
| pdx1 | ENSAMXG00000031179 | GATCTCTGTCCAGAGCGAAAC | CGCTTGTTGTCCTCCACTT |
| per2 | ENSAMXG00000001431 | CGTCATCCAGCGAGAGTAAC | TCCTGAAAGGTTCTAGGTTTGG |
| ghrl | ENSAMXG00000002173 | TTACTTGTGGCTCCAGCTTC | AAGGTGCGCTCATCATTAGG |
| nr1d1 | ENSAMXG00000015855 | GGAACCTTTGAGGTGCTGAT | GTAGGTGGTGCCAGAGATAAAC |
| deptor | ENSAMXG00000042482 | GCTGTTGGAATGAAGGTGTG | TGAGGATGAGGTGACTGACT |
| duox | ENSAMXG00000000686 | ACGTCTTTGCTACGCACTAC | GGATCACCGTCAAGGTGTAAA |
| fads2 | ENSAMXG00000015974 | CATGCTGAAGATCCTGGTACTG | GCAGAGGAGGACCAATGAAA |
| dnmt3bb.1 | ENSAMXG00000019567 | TCAGCCAATGCTGTCATACG | TGGGCATTCAAACTCCTGTC |
| dnmt1 | ENSAMXG00000012182 | CGTCCGGAACTTTGTGTCTTT | ACTGTCCAGCCTGAAGTACA |
| tulp1a | ENSAMXG00000005179 | GTTCACAGCAAAGACAGTGATTAT | CGCACAGCGGGTACTTATAG |
| mob4 | ENSAMXG00000036359 | TCTGGCAGTGCAACAGTATATC | CGTACTTCCACACTCCTTCATC |
| tbcb | ENSAMXG00000038054 | TTGTTCAGCACCTCGGATAAG | CGGTCGATCACGTGTATTCTG |
| rnf7 | ENSAMXG00000029464 | GTCATGGATGCGTGTCTGA | GACATGCAGCAGTTATGGAAAG |

Supplementary Table 2. Summary statistics of Illumina output: the number reads, total base pairs, quality trimmed reads retained for each treatment, and the overall mapping rate from Tophat2 using Bowtie2.

| Treatment | Illumina reads | Total Base Pairs | Clean Reads | Overall Mapping Rate (%) |
|---|---|---|---|---|
| Dark 1 | 24649247 | 7394774100 | 38834866 | 67.7 |
| Dark 2 | 21923603 | 6577080900 | 33789396 | 68.8 |
| Dark 3 | 24983994 | 7495198200 | 39617512 | 72.9 |
| Light 1 | 27659128 | 8297738400 | 41958430 | 60.0 |
| Light 2 | 23643113 | 7092933900 | 36800490 | 57.6 |
| Light 3 | 25549410 | 7664823000 | 39436392 | 61.8 |

Supplementary Table 3. List of differentially expressed genes (DEG) between three surface fish (7 month-old) placed in either totally dark (D/D) or light/ dark (L/D) conditions within 1 day post fertilization. Differential expression was determined using the Cufflinks pipeline. Significance threshold was set at $p_{adj}<0.1$.

| Locus | gene_short_name | Dark(FPKM) | Light(FPKM) | Light_Dark_p_value | Light_Dark_q_value | Light_Dark_value_1 | Light_Dark_value_2 | Light_Dark_log2_fold_change | Light_Dark_test_stat | length |
|---|---|---|---|---|---|---|---|---|---|---|
| XLOC_000025 | sytl5 | 1.72938 | 4.90491 | 1e-04 | 0.0124776 | 1.72938 | 4.90491 | 1.50397 | 2.20332 | KB871578.1:1440714-1462997 |
| XLOC_000093 | inhbb | 2.81404 | 10.2465 | 5e-05 | 0.00738566 | 2.81404 | 10.2465 | 1.86442 | 2.82349 | KB871578.1:7510497-7518815 |
| XLOC_000111 | gpr143 | 2.81896 | 0.994148 | 0.0013 | 0.0811047 | 2.81896 | 0.994148 | -1.50363 | -1.92644 | KB871578.1:492654-518016 |
| XLOC_000262 | rag1 | 0.888883 | 3.23342 | 0.00045 | 0.0403574 | 0.888883 | 3.23342 | 1.86299 | 2.17165 | KB871579.1:5211102-5217994 |
| XLOC_000295 | nr2e3 | 6.71822 | 1.71059 | 5e-05 | 0.00738566 | 6.71822 | 1.71059 | -1.97358 | -2.89312 | KB871579.1:8814852-8819179 |
| XLOC_000399 | actc1c | 23.6314 | 9.25617 | 0.00075 | 0.0540027 | 23.6314 | 9.25617 | -1.35222 | -1.98907 | KB871580.1:16121-28370 |
| XLOC_000433 | asb12b | 8.43801 | 3.26271 | 0.00085 | 0.0574938 | 8.43801 | 3.26271 | -1.37083 | -1.95768 | KB871580.1:3001595-3006229 |
| XLOC_000452 | zgc:174917 | 8.95196 | 57.3576 | 5e-05 | 0.00738566 | 8.95196 | 57.3576 | 2.67971 | 3.59156 | KB871580.1:5079647-5085941 |
| XLOC_000453 | si:ch211-153b23.4 | 11.8002 | 73.0788 | 5e-05 | 0.00738566 | 11.8002 | 73.0788 | 2.63065 | 3.34646 | KB871580.1:5095627-5104246 |
| XLOC_000454 | irg1l | 22.5864 | 132.136 | 5e-05 | 0.00738566 | 22.5864 | 132.136 | 2.54849 | 2.91901 | KB871580.1:5113417-5116993 |
| XLOC_000485 | dyrk4 | 11.7941 | 3.88336 | 5e-05 | 0.00738566 | 11.7941 | 3.88336 | -1.60269 | -2.4286 | KB871580.1:2111547-2129387 |
| XLOC_000515 | si:ch211-153b23.5 | 157.673 | 712.965 | 0.00055 | 0.0450978 | 157.673 | 712.965 | 2.1769 | 2.27869 | KB871580.1:5088174-5094399 |
| XLOC_000580 | golt1a | 12.1484 | 3.97679 | 5e-05 | 0.00738566 | 12.1484 | 3.97679 | -1.6111 | -2.3119 | KB871584.1:999779-1004564 |
| XLOC_000649 | rbp2b | 2.60755 | 0.53257 | 0.001 | 0.0665199 | 2.60755 | 0.53257 | -2.29165 | -2.18177 | KB871588.1:322775-327198 |

| Locus | gene_short_name | Dark(FPKM) | Light(FPKM) | Light_Dark_p_value | Light_Dark_q_value | Light_Dark_value_1 | Light_Dark_value_2 | Light_Dark_log2_fold_change | Light_Dark_test_stat | length |
|---|---|---|---|---|---|---|---|---|---|---|
| XLOC_000702 | ENSAMXG00000016396 | 15.7313 | 6.56485 | 0.0015 | 0.0878528 | 15.7313 | 6.56485 | -1.2608 | -1.87407 | KB871590.1:710199-717388 |
| XLOC_000735 | ENSAMXG00000002944 | 0.635797 | 3.46134 | 7e-04 | 0.0518904 | 0.635797 | 3.46134 | 2.44469 | 2.15452 | KB871592.1:79745-81396 |
| XLOC_000771 | mpx | 17.9612 | 58.2451 | 5e-05 | 0.00738566 | 17.9612 | 58.2451 | 1.69725 | 2.58898 | KB871593.1:122277-151875 |
| XLOC_000852 | ftcd | 8.39321 | 3.25075 | 5e-05 | 0.00738566 | 8.39321 | 3.25075 | -1.36845 | -2.12144 | KB871596.1:296241-307344 |
| XLOC_000916 | ENSAMXG00000008669 | 2.28762 | 8.78731 | 0.00055 | 0.0450978 | 2.28762 | 8.78731 | 1.94158 | 2.31209 | KB871598.1:979360-1015819 |
| XLOC_000921 | tecra | 19.2998 | 8.25754 | 0.00155 | 0.0902549 | 19.2998 | 8.25754 | -1.2248 | -1.89801 | KB871599.1:37705-47301 |
| XLOC_000952 | gcgra | 1.08799 | 2.77062 | 0.00125 | 0.0789662 | 1.08799 | 2.77062 | 1.34855 | 1.89347 | KB871599.1:242400-262385 |
| XLOC_000970 | soul5 | 9.69748 | 44.2789 | 5e-05 | 0.00738566 | 9.69748 | 44.2789 | 2.19094 | 2.8049 | KB871599.1:974982-1005025 |
| XLOC_000984 | unc45b | 29.1538 | 8.21331 | 5e-05 | 0.00738566 | 29.1538 | 8.21331 | -1.82765 | -2.7806 | KB871601.1:371258-381174 |
| XLOC_001030 | ENSAMXG00000013132 | 3.5872 | 1.23813 | 0.00145 | 0.0861806 | 3.5872 | 1.23813 | -1.53469 | -1.91854 | KB871601.1:1187756-1192073 |
| XLOC_001093 | si:ch211-244a23.1 | 4.09304 | 1.05886 | 5e-05 | 0.00738566 | 4.09304 | 1.05886 | -1.95066 | -2.68567 | KB871603.1:346775-425210 |
| XLOC_001264 | nr4a3 | 1.48495 | 3.71457 | 5e-04 | 0.0432953 | 1.48495 | 3.71457 | 1.32278 | 2.02312 | KB871610.1:518753-561832 |
| XLOC_001293 | rln3b | 3.74295 | 0.853494 | 0.00075 | 0.0540027 | 3.74295 | 0.853494 | -2.13272 | -2.49295 | KB871612.1:143776-147078 |
| XLOC_001328 | crtac1a | 2.13533 | 11.401 | 5e-05 | 0.00738566 | 2.13533 | 11.401 | 2.41662 | 3.5946 | KB871612.1:859389-870145 |
| XLOC_001370 | pmela | 8.66005 | 3.77339 | 0.0012 | 0.0770185 | 8.66005 | 3.77339 | -1.19852 | -1.86439 | KB871614.1:1074424-1091771 |
| XLOC_001576 | itga6l | 6.81959 | 33.3774 | 5e-05 | 0.00738566 | 6.81959 | 33.3774 | 2.29111 | 3.2917 | KB871621.1:913994-946351 |

| Locus | gene_short_name | Dark(FPKM) | Light(FPKM) | Light_Dark_p_value | Light_Dark_q_value | Light_Dark_value_1 | Light_Dark_value_2 | Light_Dark_log2_fold_change | Light_Dark_test_stat | length |
|---|---|---|---|---|---|---|---|---|---|---|
| XLOC_001598 | tlr5a | 1.58578 | 4.00942 | 0.00145 | 0.0861806 | 1.58578 | 4.00942 | 1.3382 | 1.90087 | KB871623.1:354083-356732 |
| XLOC_001630 | ENSAMXG00000012407 | 7.88452 | 0.972725 | 5e-05 | 0.00738566 | 7.88452 | 0.972725 | -3.01892 | -3.58831 | KB871624.1:632513-689013 |
| XLOC_001660 | ENSAMXG00000015910 | 0.677565 | 3.24349 | 5e-05 | 0.00738566 | 0.677565 | 3.24349 | 2.25912 | 2.56101 | KB871626.1:91983-98799 |
| XLOC_001783 | ENSAMXG00000021817 | 4.54948 | 1.27881 | 2e-04 | 0.0216011 | 4.54948 | 1.27881 | -1.8309 | -2.3516 | KB871630.1:5861922-5865492 |
| XLOC_001824 | tyrp1b | 1.50872 | 0.276749 | 2e-04 | 0.0216011 | 1.50872 | 0.276749 | -2.44668 | -2.7465 | KB871630.1:5217534-5227365 |
| XLOC_001840 | fgg | 61.5504 | 21.1568 | 3e-04 | 0.0296882 | 61.5504 | 21.1568 | -1.54065 | -2.08075 | KB871630.1:5844873-5851876 |
| XLOC_001917 | ENSAMXG00000028266 | 32.4304 | 10.5147 | 5e-05 | 0.00738566 | 32.4304 | 10.5147 | -1.62494 | -2.56256 | KB871633.1:1064227-1073679 |
| XLOC_001944 | ENSAMXG00000003102 | 90.7779 | 790.256 | 5e-05 | 0.00738566 | 90.7779 | 790.256 | 3.12191 | 3.2252 | KB871635.1:35154-42471 |
| XLOC_001955 | obsl1b | 7.83515 | 3.47243 | 0.00115 | 0.0742841 | 7.83515 | 3.47243 | -1.17401 | -1.81193 | KB871635.1:731204-770758 |
| XLOC_001973 | lipca | 2.1924 | 0.167918 | 0.00105 | 0.0689328 | 2.1924 | 0.167918 | -3.70668 | -2.88502 | KB871636.1:533700-560666 |
| XLOC_002044 | si:ch211-81a5.8 | 6.41447 | 0.866872 | 5e-05 | 0.00738566 | 6.41447 | 0.866872 | -2.88744 | -3.42552 | KB871640.1:869861-877544 |
| XLOC_002047 | ENSAMXG00000010135 | 87.6827 | 22.9572 | 5e-05 | 0.00738566 | 87.6827 | 22.9572 | -1.93334 | -2.52617 | KB871640.1:1055303-1099497 |
| XLOC_002080 | cuzd1.2 | 64.7338 | 19.8308 | 5e-05 | 0.00738566 | 64.7338 | 19.8308 | -1.70678 | -2.69252 | KB871643.1:701902-721166 |
| XLOC_002086 | htra1a | 7.4593 | 3.14847 | 0.00075 | 0.0540027 | 7.4593 | 3.14847 | -1.24439 | -1.95411 | KB871643.1:547526-613060 |
| XLOC_002112 | ENSAMXG00000004676 | 56.1736 | 25.2781 | 0.0013 | 0.0811047 | 56.1736 | 25.2781 | -1.15201 | -1.89297 | KB871644.1:889063-897015 |
| XLOC_002320 | ENSAMXG00000009532 | 6.6181 | 16.2928 | 8e-04 | 0.0556097 | 6.6181 | 16.2928 | 1.29975 | 1.99253 | KB871654.1:944058-952616 |

| Locus | gene_short_name | Dark(FPKM) | Light(FPKM) | Light_Dark_p_value | Light_Dark_q_value | Light_Dark_value_1 | Light_Dark_value_2 | Light_Dark_log2_fold_change | Light_Dark_test_stat | length |
|---|---|---|---|---|---|---|---|---|---|---|
| XLOC_002453 | ENSAMXG00000001841 | 8.72091 | 3.03604 | 0.00075 | 0.0540027 | 8.72091 | 3.03604 | -1.52229 | -2.00856 | KB871662.1:842321-847168 |
| XLOC_002460 | apoa2 | 1446.47 | 417.417 | 2e-04 | 0.0216011 | 1446.47 | 417.417 | -1.79297 | -2.10386 | KB871662.1:755634-759753 |
| XLOC_002461 | apoc4 | 13.0281 | 4.64507 | 0.0016 | 0.0923632 | 13.0281 | 4.64507 | -1.48786 | -1.95224 | KB871662.1:763921-766276 |
| XLOC_002528 | ENSAMXG00000015132 | 11.122 | 32.2114 | 5e-05 | 0.00738566 | 11.122 | 32.2114 | 1.53415 | 2.31617 | KB871666.1:617191-660944 |
| XLOC_002544 | ENSAMXG00000018298 | 1.48336 | 0 | 5e-05 | 0.00738566 | 1.48336 | 0 | -Inf | NA | KB871667.1:715875-729135 |
| XLOC_002557 | casq1b | 39.448 | 12.6542 | 1e-04 | 0.0124776 | 39.448 | 12.6542 | -1.64034 | -2.51524 | KB871667.1:596086-630314 |
| XLOC_002592 | pde6ha | 153.962 | 990.1 | 5e-05 | 0.00738566 | 153.962 | 990.1 | 2.685 | 4.25525 | KB871670.1:579803-580207 |
| XLOC_002599 | si:zfos-323e3.4 | 2.33197 | 6.29524 | 8e-04 | 0.0556097 | 2.33197 | 6.29524 | 1.43271 | 2.09449 | KB871670.1:218416-232095 |
| XLOC_002665 | ENSAMXG00000007037 | 1524.76 | 509.863 | 4e-04 | 0.0372019 | 1524.76 | 509.863 | -1.5804 | -2.06446 | KB871673.1:828202-829527 |
| XLOC_002845 | ENSAMXG00000015374 | 26.3848 | 11.1454 | 0.00085 | 0.0574938 | 26.3848 | 11.1454 | -1.24326 | -1.92956 | KB871682.1:405661-408875 |
| XLOC_002905 | ENSAMXG00000017046 | 18.7434 | 101.724 | 5e-05 | 0.00738566 | 18.7434 | 101.724 | 2.44021 | 2.84071 | KB871685.1:170515-179715 |
| XLOC_003111 | ENSAMXG00000007743 | 489.302 | 3068.49 | 5e-05 | 0.00738566 | 489.302 | 3068.49 | 2.64873 | 2.73331 | KB871688.1:5644019-5646112 |
| XLOC_003118 | nt5c2l1 | 1.31007 | 3.22673 | 0.00085 | 0.0574938 | 1.31007 | 3.22673 | 1.30043 | 1.91801 | KB871689.1:45593-61053 |
| XLOC_003124 | slc5a1 | 13.5913 | 47.7812 | 5e-05 | 0.00738566 | 13.5913 | 47.7812 | 1.81376 | 2.89548 | KB871689.1:411283-432176 |
| XLOC_003129 | ENSAMXG00000028718 | 15.9252 | 63.1131 | 5e-05 | 0.00738566 | 15.9252 | 63.1131 | 1.98663 | 3.07609 | KB871689.1:749657-763337 |
| XLOC_003159 | aglb | 8.2341 | 3.12566 | 1e-04 | 0.0124776 | 8.2341 | 3.12566 | -1.39745 | -2.25824 | KB871691.1:425429-466515 |

| Locus | gene_short_name | Dark(FPKM) | Light(FPKM) | Light_Dark_p_value | Light_Dark_q_value | Light_Dark_value_1 | Light_Dark_value_2 | Light_Dark_log2_fold_change | Light_Dark_test_stat | length |
|---|---|---|---|---|---|---|---|---|---|---|
| XLOC_003208 | ENSAMXG00000014187 | 26.4581 | 7.27278 | 5e-05 | 0.00738566 | 26.4581 | 7.27278 | -1.86313 | -2.39019 | KB871694.1:716826-718209 |
| XLOC_003273 | srl | 14.8543 | 5.28934 | 5e-05 | 0.00738566 | 14.8543 | 5.28934 | -1.48972 | -2.33184 | KB871702.1:150751-170936 |
| XLOC_003307 | nr1d1 | 13.821 | 5.389 | 5e-05 | 0.00738566 | 13.821 | 5.389 | -1.35877 | -2.20071 | KB871704.1:426428-435796 |
| XLOC_003375 | ENSAMXG00000014747 | 73.4764 | 26.5773 | 0.00085 | 0.0574938 | 73.4764 | 26.5773 | -1.46709 | -1.98389 | KB871707.1:35060-44134 |
| XLOC_003531 | cyp8b2 | 17.2369 | 5.96801 | 5e-05 | 0.00738566 | 17.2369 | 5.96801 | -1.53018 | -2.28808 | KB871717.1:704847-706458 |
| XLOC_003553 | ENSAMXG00000014372 | 4.50593 | 1.48769 | 5e-04 | 0.0432953 | 4.50593 | 1.48769 | -1.59874 | -2.1852 | KB871718.1:576394-607652 |
| XLOC_003924 | mybpha | 39.8948 | 14.4617 | 0.00015 | 0.0178305 | 39.8948 | 14.4617 | -1.46396 | -2.36596 | KB871745.1:680467-705649 |
| XLOC_003980 | maats1 | 1.25126 | 0.439702 | 0.00125 | 0.0789662 | 1.25126 | 0.439702 | -1.50878 | -1.95201 | KB871750.1:479109-515783 |
| XLOC_003987 | gpx7 | 5.53336 | 1.77059 | 1e-04 | 0.0124776 | 5.53336 | 1.77059 | -1.64392 | -2.22967 | KB871751.1:456402-470647 |
| XLOC_004077 | uox | 25.9104 | 8.36595 | 5e-05 | 0.00738566 | 25.9104 | 8.36595 | -1.63093 | -2.36451 | KB871753.1:1074353-1080629 |
| XLOC_004096 | ENSAMXG00000017658 | 25.8306 | 8.66116 | 0.00045 | 0.0403574 | 25.8306 | 8.66116 | -1.57645 | -2.08683 | KB871753.1:2178555-2200020 |
| XLOC_004174 | tgm2a | 17.5466 | 3.81582 | 5e-05 | 0.00738566 | 17.5466 | 3.81582 | -2.20113 | -3.14917 | KB871754.1:5123302-5130631 |
| XLOC_004261 | sst1.2 | 15.8065 | 109.086 | 5e-05 | 0.00738566 | 15.8065 | 109.086 | 2.78688 | 3.1029 | KB871758.1:283368-286701 |
| XLOC_004289 | ENSAMXG00000002805 | 5.04307 | 28.0732 | 5e-05 | 0.00738566 | 5.04307 | 28.0732 | 2.47682 | 3.45563 | KB871761.1:16057-18860 |
| XLOC_004290 | ENSAMXG00000002840 | 0.400355 | 2.39441 | 3e-04 | 0.0296882 | 0.400355 | 2.39441 | 2.58032 | 2.6615 | KB871761.1:23088-33199 |
| XLOC_004309 | rbp3 | 19.536 | 55.9164 | 5e-05 | 0.00738566 | 19.536 | 55.9164 | 1.51714 | 2.42609 | KB871761.1:382801-385067 |

| Locus | gene_short_name | Dark(FPKM) | Light(FPKM) | Light_Dark_p_value | Light_Dark_q_value | Light_Dark_value_1 | Light_Dark_value_2 | Light_Dark_log2_fold_change | Light_Dark_test_stat | length |
|---|---|---|---|---|---|---|---|---|---|---|
| XLOC_004337 | ENSAMXG00000018234 | 6.72252 | 2.51078 | 0.00055 | 0.0450978 | 6.72252 | 2.51078 | -1.42087 | -2.00482 | KB871763.1:444998-472622 |
| XLOC_004372 | col10a1a | 125.49 | 41.3909 | 1e-04 | 0.0124776 | 125.49 | 41.3909 | -1.60019 | -2.24698 | KB871767.1:433097-441907 |
| XLOC_004484 | txn | 165.002 | 359.659 | 0.0015 | 0.0878528 | 165.002 | 359.659 | 1.12415 | 1.79726 | KB871773.1:351811-358272 |
| XLOC_004868 | ENSAMXG00000004646 | 4.34645 | 1.54864 | 0.00055 | 0.0450978 | 4.34645 | 1.54864 | -1.48883 | -2.0516 | KB871797.1:262399-274704 |
| XLOC_004873 | tm4sf4 | 530.68 | 162.437 | 0.00025 | 0.0254937 | 530.68 | 162.437 | -1.70797 | -2.29356 | KB871797.1:586846-596495 |
| XLOC_004879 | ENSAMXG00000016751 | 2.14144 | 5.73489 | 0.00025 | 0.0254937 | 2.14144 | 5.73489 | 1.42118 | 2.09893 | KB871798.1:173791-194682 |
| XLOC_004908 | ENSAMXG00000013747 | 1.84443 | 7.24206 | 5e-04 | 0.0432953 | 1.84443 | 7.24206 | 1.97322 | 2.24199 | KB871800.1:304609-308211 |
| XLOC_004996 | ENSAMXG00000013876 | 1.5781 | 9.6664 | 2e-04 | 0.0216011 | 1.5781 | 9.6664 | 2.61479 | 2.46262 | KB871804.1:568886-571789 |
| XLOC_005033 | ENSAMXG00000007580 | 7.63171 | 2.40503 | 5e-05 | 0.00738566 | 7.63171 | 2.40503 | -1.66595 | -2.48256 | KB871807.1:217452-235079 |
| XLOC_005217 | fabp10a | 513.26 | 122.233 | 5e-05 | 0.00738566 | 513.26 | 122.233 | -2.07005 | -2.62509 | KB871824.1:186324-193248 |
| XLOC_005233 | rorca | 3.93054 | 10.2191 | 2e-04 | 0.0216011 | 3.93054 | 10.2191 | 1.37847 | 2.07223 | KB871825.1:273511-307601 |
| XLOC_005239 | nexn | 17.6518 | 7.06033 | 6e-04 | 0.0484072 | 17.6518 | 7.06033 | -1.32201 | -2.03959 | KB871825.1:551365-582477 |
| XLOC_005272 | cps1 | 3.07537 | 1.26253 | 0.00055 | 0.0450978 | 3.07537 | 1.26253 | -1.28445 | -1.87905 | KB871827.1:115446-246773 |
| XLOC_005342 | hmox1a | 1.63559 | 7.73547 | 5e-05 | 0.00738566 | 1.63559 | 7.73547 | 2.24168 | 2.75381 | KB871831.1:280111-282576 |
| XLOC_005499 | fads2 | 142.981 | 36.8543 | 5e-05 | 0.00738566 | 142.981 | 36.8543 | -1.95592 | -2.92632 | KB871834.1:3460641-3478702 |
| XLOC_005516 | hal | 9.97762 | 3.83312 | 5e-04 | 0.0432953 | 9.97762 | 3.83312 | -1.38018 | -2.01239 | KB871834.1:4355479-4371595 |

| Locus | gene_short_name | Dark(FPKM) | Light(FPKM) | Light_Dark_p_value | Light_Dark_q_value | Light_Dark_value_1 | Light_Dark_value_2 | Light_Dark_log2_fold_change | Light_Dark_test_stat | length |
|---|---|---|---|---|---|---|---|---|---|---|
| XLOC_005520 | tph1a | 13.4651 | 4.63457 | 1e-04 | 0.0124776 | 13.4651 | 4.63457 | -1.53872 | -2.27977 | KB871834.1:4504607-4511760 |
| XLOC_005569 | ENSAMXG00000004797 | 267.793 | 80.7565 | 0.00025 | 0.0254937 | 267.793 | 80.7565 | -1.72947 | -2.10168 | KB871837.1:315052-319078 |
| XLOC_005599 | clul1 | 5.57972 | 12.8579 | 0.00145 | 0.0861806 | 5.57972 | 12.8579 | 1.20439 | 1.82359 | KB871840.1:280159-290652 |
| XLOC_005798 | per2 | 2.00363 | 6.60999 | 5e-05 | 0.00738566 | 2.00363 | 6.60999 | 1.72203 | 2.71497 | KB871856.1:269900-309108 |
| XLOC_005825 | gpbar1 | 1.19921 | 3.67477 | 0.0014 | 0.0852261 | 1.19921 | 3.67477 | 1.61557 | 1.97244 | KB871858.1:599602-600616 |
| XLOC_005958 | nr1d4a | 3.49854 | 1.3138 | 0.00105 | 0.0689328 | 3.49854 | 1.3138 | -1.41301 | -1.89739 | KB871870.1:467456-493980 |
| XLOC_005996 | slc39a8 | 11.5426 | 4.6383 | 9e-04 | 0.0606715 | 11.5426 | 4.6383 | -1.31531 | -1.86312 | KB871873.1:449404-454954 |
| XLOC_006012 | nmbb | 2.41993 | 0 | 5e-05 | 0.00738566 | 2.41993 | 0 | -Inf | NA | KB871875.1:56-661 |
| XLOC_006122 | zgc:85843 | 24.0139 | 8.05679 | 1e-04 | 0.0124776 | 24.0139 | 8.05679 | -1.57559 | -2.25041 | KB871883.1:60492-87189 |
| XLOC_006155 | ENSAMXG00000012871 | 16.6802 | 5.64746 | 1e-04 | 0.0124776 | 16.6802 | 5.64746 | -1.56246 | -2.43038 | KB871886.1:202267-238321 |
| XLOC_006189 | kdm6bb | 7.10864 | 2.55495 | 0.0016 | 0.0923632 | 7.10864 | 2.55495 | -1.47628 | -1.96655 | KB871888.1:152503-181265 |
| XLOC_006192 | hhatla | 58.8582 | 23.0381 | 4e-04 | 0.0372019 | 58.8582 | 23.0381 | -1.35322 | -2.11597 | KB871889.1:2464-18923 |
| XLOC_006425 | ENSAMXG00000012617 | 23.952 | 8.70033 | 2e-04 | 0.0216011 | 23.952 | 8.70033 | -1.461 | -2.09588 | KB871909.1:523073-526998 |
| XLOC_006426 | ENSAMXG00000012633 | 19.7979 | 7.31712 | 0.00065 | 0.0502225 | 19.7979 | 7.31712 | -1.436 | -1.9465 | KB871909.1:531694-533828 |
| XLOC_006545 | krtt1c19e | 24.6566 | 8.80509 | 0.00085 | 0.0574938 | 24.6566 | 8.80509 | -1.48556 | -2.11358 | KB871921.1:197020-204202 |
| XLOC_006633 | atp1a2a | 12.584 | 3.79992 | 5e-05 | 0.00738566 | 12.584 | 3.79992 | -1.72754 | -2.709 | KB871931.1:154312-213455 |

| Locus | gene_short_name | Dark(FPKM) | Light(FPKM) | Light_Dark_p_value | Light_Dark_q_value | Light_Dark_value_1 | Light_Dark_value_2 | Light_Dark_log2_fold_change | Light_Dark_test_stat | length |
|---|---|---|---|---|---|---|---|---|---|---|
| XLOC_006714 | guca1c | 3.47989 | 10.9861 | 5e-05 | 0.00738566 | 3.47989 | 10.9861 | 1.65856 | 2.41026 | KB871938.1:1023233-1030654 |
| XLOC_006715 | thrsp | 138.457 | 14.6556 | 5e-05 | 0.00738566 | 138.457 | 14.6556 | -3.23991 | -3.87353 | KB871938.1:1035629-1036046 |
| XLOC_006754 | ENSAMXG00000010569 | 81.7739 | 16.2647 | 0.00045 | 0.0403574 | 81.7739 | 16.2647 | -2.3299 | -2.1627 | KB871938.1:4970804-5088665 |
| XLOC_006824 | lonrf1l | 2.60428 | 7.09716 | 0.00015 | 0.0178305 | 2.60428 | 7.09716 | 1.44636 | 2.25755 | KB871939.1:998737-1008850 |
| XLOC_006867 | slc44a5b | 3.0143 | 8.15188 | 7e-04 | 0.0518904 | 3.0143 | 8.15188 | 1.43531 | 1.99033 | KB871939.1:112534-168315 |
| XLOC_006902 | ENSAMXG00000014892 | 1.15661 | 3.16266 | 1e-04 | 0.0124776 | 1.15661 | 3.16266 | 1.45123 | 2.28706 | KB871939.1:2448361-2495470 |
| XLOC_007020 | ENSAMXG00000014157 | 8.75821 | 0 | 2e-04 | 0.0216011 | 8.75821 | 0 | -Inf | NA | KB871956.1:282358-292207 |
| XLOC_007022 | cers3a | 3.20073 | 8.61021 | 0.00025 | 0.0254937 | 3.20073 | 8.61021 | 1.42765 | 2.17536 | KB871956.1:344505-378639 |
| XLOC_007029 | ENSAMXG00000004919 | 2.34166 | 7.6518 | 0.00015 | 0.0178305 | 2.34166 | 7.6518 | 1.70827 | 2.26338 | KB871957.1:267374-285774 |
| XLOC_007097 | si:ch73-366l1.5 | 31.1437 | 8.53223 | 5e-05 | 0.00738566 | 31.1437 | 8.53223 | -1.86795 | -2.84312 | KB871968.1:178450-191470 |
| XLOC_007139 | ENSAMXG00000017732 | 18.9833 | 7.30933 | 7e-04 | 0.0518904 | 18.9833 | 7.30933 | -1.37692 | -2.02268 | KB871971.1:44926-183149 |
| XLOC_007244 | acsl3a | 1.30106 | 3.53938 | 0.00035 | 0.0338036 | 1.30106 | 3.53938 | 1.44381 | 2.01865 | KB871983.1:414440-478193 |
| XLOC_007276 | opn1sw2 | 18.629 | 50.3663 | 5e-05 | 0.00738566 | 18.629 | 50.3663 | 1.43491 | 2.31712 | KB871987.1:241053-248021 |
| XLOC_007454 | hjv | 6.36292 | 1.99583 | 5e-05 | 0.00738566 | 6.36292 | 1.99583 | -1.6727 | -2.47429 | KB872011.1:25030-30978 |
| XLOC_007827 | ENSAMXG00000016413 | 18.8718 | 4.65029 | 5e-05 | 0.00738566 | 18.8718 | 4.65029 | -2.02084 | -2.40919 | KB872051.1:117984-119391 |
| XLOC_007828 | ENSAMXG00000016426 | 13.2634 | 2.54315 | 5e-05 | 0.00738566 | 13.2634 | 2.54315 | -2.38277 | -2.58315 | KB872051.1:128145-129335 |

| Locus | gene_short_name | Dark(FPKM) | Light(FPKM) | Light_Dark_p_value | Light_Dark_q_value | Light_Dark_value_1 | Light_Dark_value_2 | Light_Dark_log2_fold_change | Light_Dark_test_stat | length |
|---|---|---|---|---|---|---|---|---|---|---|
| XLOC_007831 | hbaa2 | 4.63625 | 14.6156 | 0.00105 | 0.0689328 | 4.63625 | 14.6156 | 1.65648 | 1.95051 | KB872051.1:192831-193838 |
| XLOC_007843 | ENSAMXG00000027508 | 15.1346 | 3.59669 | 5e-05 | 0.00738566 | 15.1346 | 3.59669 | -2.07311 | -2.57358 | KB872053.1:242757-247769 |
| XLOC_007871 | ENSAMXG00000014852 | 1.46566 | 4.56649 | 0.00115 | 0.0742841 | 1.46566 | 4.56649 | 1.63954 | 2.03722 | KB872056.1:223244-234328 |
| XLOC_007938 | lims1 | 11.6389 | 4.71026 | 0.00035 | 0.0338036 | 11.6389 | 4.71026 | -1.30508 | -1.98877 | KB872067.1:60303-75465 |
| XLOC_007949 | ENSAMXG00000011973 | 4.27761 | 0.542253 | 0.00175 | 0.0987521 | 4.27761 | 0.542253 | -2.97977 | -2.67761 | KB872068.1:62057-64354 |
| XLOC_008008 | ENSAMXG00000002825 | 84.4957 | 35.7196 | 0.00145 | 0.0861806 | 84.4957 | 35.7196 | -1.24216 | -1.93292 | KB872073.1:67872-84597 |
| XLOC_008156 | oit3 | 18.2981 | 8.02205 | 0.00155 | 0.0902549 | 18.2981 | 8.02205 | -1.18965 | -1.84166 | KB872081.1:3943983-3962613 |
| XLOC_008165 | ENSAMXG00000004102 | 36.2807 | 12.7707 | 4e-04 | 0.0372019 | 36.2807 | 12.7707 | -1.50636 | -2.08899 | KB872081.1:5025369-5034203 |
| XLOC_008312 | ENSAMXG00000005092 | 160.199 | 53.533 | 2e-04 | 0.0216011 | 160.199 | 53.533 | -1.58136 | -2.221 | KB872099.1:5050-16559 |
| XLOC_008418 | ENSAMXG00000001148 | 68.0436 | 15.6074 | 5e-05 | 0.00738566 | 68.0436 | 15.6074 | -2.12422 | -3.14949 | KB872110.1:134175-138156 |
| XLOC_008522 | fgfbp2a | 14.1128 | 32.843 | 0.00085 | 0.0574938 | 14.1128 | 32.843 | 1.21859 | 1.94405 | KB872123.1:87872-90425 |
| XLOC_008553 | arr3a | 13.9001 | 58.9019 | 5e-05 | 0.00738566 | 13.9001 | 58.9019 | 2.08322 | 3.07692 | KB872126.1:165767-172954 |
| XLOC_008650 | ENSAMXG00000010829 | 5.18706 | 1.51229 | 6e-04 | 0.0484072 | 5.18706 | 1.51229 | -1.77818 | -2.20244 | KB872142.1:228352-245718 |
| XLOC_008843 | apoa1b | 1185.73 | 290.254 | 8e-04 | 0.0556097 | 1185.73 | 290.254 | -2.0304 | -2.21997 | KB872172.1:161545-162549 |
| XLOC_009073 | slc34a2a | 4.59594 | 1.03457 | 5e-05 | 0.00738566 | 4.59594 | 1.03457 | -2.15133 | -2.72931 | KB872220.1:103546-112146 |
| XLOC_009276 | ENSAMXG00000002779 | 3.77377 | 13.5332 | 2e-04 | 0.0216011 | 3.77377 | 13.5332 | 1.84242 | 2.52683 | KB872260.1:223227-232193 |

| Locus | gene_short_name | Dark(FPKM) | Light(FPKM) | Light_Dark_p_value | Light_Dark_q_value | Light_Dark_value_1 | Light_Dark_value_2 | Light_Dark_log2_fold_change | Light_Dark_test_stat | length |
|---|---|---|---|---|---|---|---|---|---|---|
| XLOC_009284 | ENSAMXG00000012870 | 6.91874 | 0 | 5e-05 | 0.00738566 | 6.91874 | 0 | -Inf | NA | KB872262.1:55697-59406 |
| XLOC_009374 | ENSAMXG00000004416 | 3.32707 | 0.709329 | 5e-05 | 0.00738566 | 3.32707 | 0.709329 | -2.22972 | -3.09356 | KB872286.1:83633-164522 |
| XLOC_009400 | ENSAMXG00000004721 | 16.0445 | 7.12806 | 0.00175 | 0.0987521 | 16.0445 | 7.12806 | -1.1705 | -1.76127 | KB872293.1:20-20286 |
| XLOC_009443 | abcb11b | 6.71758 | 0.910457 | 5e-05 | 0.00738566 | 6.71758 | 0.910457 | -2.88328 | -3.91516 | KB872295.1:2665897-2687825 |
| XLOC_009522 | ENSAMXG00000018710 | 47.8519 | 21.0951 | 0.00115 | 0.0742841 | 47.8519 | 21.0951 | -1.18167 | -1.89171 | KB872295.1:2651583-2658581 |
| XLOC_009582 | pyroxd2 | 12.6538 | 4.95315 | 0.00045 | 0.0403574 | 12.6538 | 4.95315 | -1.35315 | -2.07949 | KB872296.1:3963794-3980918 |
| XLOC_009613 | ENSAMXG00000001960 | 2.73837 | 7.05334 | 6e-04 | 0.0484072 | 2.73837 | 7.05334 | 1.36499 | 2.07965 | KB872296.1:2008216-2033838 |
| XLOC_009799 | noxo1a | 2.34339 | 7.36835 | 0.00025 | 0.0254937 | 2.34339 | 7.36835 | 1.65275 | 2.26632 | KB872330.1:2661-6570 |
| XLOC_009836 | zgc:194242 | 0.421738 | 6.8048 | 5e-05 | 0.00738566 | 0.421738 | 6.8048 | 4.01213 | 3.96289 | KB872337.1:1304-8307 |
| XLOC_009839 | tulp1a | 0.723851 | 5.85016 | 5e-05 | 0.00738566 | 0.723851 | 5.85016 | 3.01471 | 4.03184 | KB872337.1:114650-123319 |
| XLOC_010145 | ENSAMXG00000027789 | 0 | 1.80021 | 5e-05 | 0.00738566 | 0 | 1.80021 | Inf | NA | KB872413.1:25255-30416 |
| XLOC_010174 | ENSAMXG00000028677 | 13.4015 | 4.46936 | 6e-04 | 0.0484072 | 13.4015 | 4.46936 | -1.58426 | -2.12762 | KB872420.1:11056-13767 |
| XLOC_010316 | ENSAMXG00000025139 | 839.084 | 239.941 | 5e-05 | 0.00738566 | 839.084 | 239.941 | -1.80613 | -2.39668 | KB872462.1:39854-40028 |
| XLOC_010317 | ENSAMXG00000016276 | 29.6083 | 9.74245 | 1e-04 | 0.0124776 | 29.6083 | 9.74245 | -1.60365 | -2.29791 | KB872462.1:45007-46706 |
| XLOC_010374 | ENSAMXG00000014782 | 83.3677 | 23.6903 | 5e-05 | 0.00738566 | 83.3677 | 23.6903 | -1.81519 | -2.59629 | KB872486.1:82927-86298 |
| XLOC_010428 | ENSAMXG00000027402 | 1.55282 | 5.57506 | 2e-04 | 0.0216011 | 1.55282 | 5.57506 | 1.8441 | 2.50389 | KB872502.1:44788-47865 |
| XLOC_010600 | nfil3-6 | 4.46653 | 10.2847 | 0.0017 | 0.0964726 | 4.46653 | 10.2847 | 1.20328 | 1.80972 | KB872561.1:31281-32529 |

| Locus | gene_short_name | Dark(FPKM) | Light(FPKM) | Light_Dark_p_value | Light_Dark_q_value | Light_Dark_value_1 | Light_Dark_value_2 | Light_Dark_log2_fold_change | Light_Dark_test_stat | length |
|---|---|---|---|---|---|---|---|---|---|---|
| XLOC_010648 | lad1 | 17.5937 | 39.3944 | 0.0017 | 0.0964726 | 17.5937 | 39.3944 | 1.16293 | 1.85973 | KB872583.1:105956-166287 |
| XLOC_010723 | ISYNA1 | 5.06362 | 12.7235 | 0.0017 | 0.0964726 | 5.06362 | 12.7235 | 1.32925 | 1.83118 | KB872617.1:42207-55231 |
| XLOC_010761 | ENSAMXG00000014761 | 5.0134 | 0.674813 | 5e-05 | 0.00738566 | 5.0134 | 0.674813 | -2.89323 | -2.86669 | KB872634.1:7919-17878 |
| XLOC_010795 | ENSAMXG00000025773 | 11.1455 | 1.18907 | 5e-05 | 0.00738566 | 11.1455 | 1.18907 | -3.22855 | -3.80008 | KB872649.1:63682-64978 |
| XLOC_010828 | ENSAMXG00000020730 | 91.073 | 27.146 | 2e-04 | 0.0216011 | 91.073 | 27.146 | -1.74628 | -2.10939 | KB872661.1:16537-18216 |
| XLOC_010858 | ENSAMXG00000000310 | 50.8205 | 21.3757 | 7e-04 | 0.0518904 | 50.8205 | 21.3757 | -1.24944 | -1.98511 | KB872672.1:53327-62597 |
| XLOC_011019 | ENSAMXG00000015089 | 9.65121 | 28.4598 | 1e-04 | 0.0124776 | 9.65121 | 28.4598 | 1.56014 | 2.44445 | KB872765.1:20868-31305 |
| XLOC_011042 | opn1mw1 | 5.61628 | 53.6355 | 5e-05 | 0.00738566 | 5.61628 | 53.6355 | 3.2555 | 4.76878 | KB872775.1:4150-8361 |
| XLOC_011065 | rorcb | 2.69209 | 6.71795 | 0.0013 | 0.0811047 | 2.69209 | 6.71795 | 1.3193 | 1.98158 | KB872789.1:22775-33957 |
| XLOC_011070 | ENSAMXG00000017636 | 131.845 | 53.3926 | 0.0014 | 0.0852261 | 131.845 | 53.3926 | -1.30413 | -1.8312 | KB872795.1:22420-23517 |
| XLOC_011143 | zgc:194930 | 4.4791 | 10.1122 | 0.00145 | 0.0861806 | 4.4791 | 10.1122 | 1.17481 | 1.87671 | KB872818.1:2447385-2478825 |
| XLOC_011258 | afp4 | 1693.33 | 542.51 | 5e-05 | 0.00738566 | 1693.33 | 542.51 | -1.64214 | -2.28476 | KB872819.1:3699762-3701692 |
| XLOC_011323 | ENSAMXG00000008228 | 2.83841 | 0.218586 | 1e-04 | 0.0124776 | 2.83841 | 0.218586 | -3.69881 | -3.65124 | KB872847.1:10343-13195 |
| XLOC_011332 | ENSAMXG00000000014 | 6.87697 | 19.705 | 0.00125 | 0.0789662 | 6.87697 | 19.705 | 1.51871 | 1.95895 | KB872850.1:43821-52867 |
| XLOC_011387 | ENSAMXG00000005597 | 79.0233 | 384.325 | 5e-05 | 0.00738566 | 79.0233 | 384.325 | 2.28198 | 2.69331 | KB872885.1:9359-15649 |
| XLOC_011414 | ENSAMXG00000002780 | 4.86323 | 1.36659 | 0.0017 | 0.0964726 | 4.86323 | 1.36659 | -1.83133 | -2.09229 | KB872914.1:11905-14091 |

| Locus | gene_short_name | Dark(FPKM) | Light(FPKM) | Light_Dark_p_value | Light_Dark_q_value | Light_Dark_value_1 | Light_Dark_value_2 | Light_Dark_log2_fold_change | Light_Dark_test_stat | length |
|---|---|---|---|---|---|---|---|---|---|---|
| XLOC_011581 | serpina7 | 14.2071 | 4.32541 | 2e-04 | 0.0216011 | 14.2071 | 4.32541 | -1.71571 | -2.33065 | KB873067.1:15187-21113 |
| XLOC_011640 | ENSAMXG00000025083 | 51.7491 | 9.08725 | 4e-04 | 0.0372019 | 51.7491 | 9.08725 | -2.50962 | -2.58218 | KB873141.1:91-1784 |
| XLOC_011673 | ENSAMXG00000015333 | 1.00377 | 4.33012 | 0.001 | 0.0665199 | 1.00377 | 4.33012 | 2.10897 | 2.11181 | KB873175.1:243-4305 |
| XLOC_011884 | fgb | 90.2955 | 24.1439 | 5e-05 | 0.00738566 | 90.2955 | 24.1439 | -1.90299 | -2.43536 | KB873510.1:1452-18168 |
| XLOC_012011 | gatm | 187.446 | 53.6218 | 5e-05 | 0.00738566 | 187.446 | 53.6218 | -1.80558 | -2.63116 | KB873821.1:646-9057 |
| XLOC_012121 | ENSAMXG00000001171 | 5.87434 | 0.608114 | 5e-05 | 0.00738566 | 5.87434 | 0.608114 | -3.27201 | -3.72064 | KB874083.1:1207-5017 |
| XLOC_012207 | ENSAMXG00000028681 | 224.641 | 88.3755 | 7e-04 | 0.0518904 | 224.641 | 88.3755 | -1.3459 | -2.03825 | KB874410.1:10781-11970 |
| XLOC_012228 | ENSAMXG00000027658 | 1.88486 | 0 | 5e-05 | 0.00738566 | 1.88486 | 0 | -Inf | NA | KB874549.1:1179-4782 |
| XLOC_012243 | socs3a | 8.69069 | 23.5974 | 0.00035 | 0.0338036 | 8.69069 | 23.5974 | 1.44109 | 2.16173 | KB874601.1:1469-2093 |
| XLOC_012247 | ENSAMXG00000007322 | 0.846368 | 7.29867 | 1e-04 | 0.0124776 | 0.846368 | 7.29867 | 3.10828 | 3.24384 | KB874626.1:2926-4177 |
| XLOC_012491 | ENSAMXG00000004196,ENSAMXG00000027240 | 12.7805 | 269.836 | 5e-05 | 0.00738566 | 12.7805 | 269.836 | 4.40006 | 9.04013 | KB876205.1:905-9614 |
| XLOC_012502 | ENSAMXG00000019672 | 21.6104 | 8.87119 | 0.00165 | 0.0947053 | 21.6104 | 8.87119 | -1.28453 | -1.84507 | KB876219.1:1225-8155 |
| XLOC_012601 | dhrs13l1 | 17.3383 | 56.2398 | 5e-05 | 0.00738566 | 17.3383 | 56.2398 | 1.69763 | 2.4985 | KB876513.1:1771-6365 |
| XLOC_012720 | RF00004 | 289.548 | 0 | 0.00135 | 0.0831906 | 289.548 | 0 | -Inf | NA | KB876961.1:5667-5785 |
| XLOC_013073 | ENSAMXG00000015463 | 1.97724 | 22.9892 | 5e-05 | 0.00738566 | 1.97724 | 22.9892 | 3.5394 | 3.75024 | KB881556.1:252-1075 |
| XLOC_013084 | plpp7 | 14.3087 | 5.64829 | 0.00025 | 0.0254937 | 14.3087 | 5.64829 | -1.34101 | -2.12568 | KB882080.1:695248-703885 |
| XLOC_013156 | ENSAMXG00000027417 | 3.30734 | 0.937835 | 0.0012 | 0.0770185 | 3.30734 | 0.937835 | -1.81827 | -1.80971 | KB882080.1:4122615-4165750 |

| Locus | gene_short_name | Dark(FPKM) | Light(FPKM) | Light_Dark_p_value | Light_Dark_q_value | Light_Dark_value_1 | Light_Dark_value_2 | Light_Dark_log2_fold_change | Light_Dark_test_stat | length |
|---|---|---|---|---|---|---|---|---|---|---|
| XLOC_013203 | si:dkey-88e18.2 | 1.38913 | 4.97042 | 8e-04 | 0.0556097 | 1.38913 | 4.97042 | 1.83918 | 1.99927 | KB882081.1:4308909-4313435 |
| XLOC_013234 | ENSAMXG00000003598 | 6.33514 | 1.44903 | 0.00035 | 0.0338036 | 6.33514 | 1.44903 | -2.12829 | -2.57682 | KB882081.1:3219975-3223639 |
| XLOC_013244 | tnni1c | 59.3591 | 24.9681 | 4e-04 | 0.0372019 | 59.3591 | 24.9681 | -1.24938 | -1.9611 | KB882081.1:4023977-4027183 |
| XLOC_013434 | obsl1a | 3.68128 | 1.59434 | 0.00065 | 0.0502225 | 3.68128 | 1.59434 | -1.20725 | -1.9129 | KB882082.1:3810391-3844914 |
| XLOC_013544 | ENSAMXG00000007167 | 14.0733 | 4.54104 | 0.00025 | 0.0254937 | 14.0733 | 4.54104 | -1.63187 | -2.21525 | KB882083.1:1898152-1908995 |
| XLOC_013638 | minpp1a | 6.28503 | 23.48 | 8e-04 | 0.0556097 | 6.28503 | 23.48 | 1.90144 | 1.96284 | KB882084.1:832325-842610 |
| XLOC_013678 | si:ch211-132f19.7 | 1.79756 | 0.474137 | 0.00025 | 0.0254937 | 1.79756 | 0.474137 | -1.92267 | -2.41254 | KB882084.1:4432035-4450238 |
| XLOC_013751 | ENSAMXG00000006428 | 1.49611 | 5.07163 | 0.00055 | 0.0450978 | 1.49611 | 5.07163 | 1.76123 | 2.08733 | KB882085.1:1710369-1713506 |
| XLOC_013762 | ENSAMXG00000006536 | 8.65892 | 29.9708 | 5e-05 | 0.00738566 | 8.65892 | 29.9708 | 1.7913 | 2.81043 | KB882085.1:2727659-2738109 |
| XLOC_013804 | f2 | 44.3624 | 15.4716 | 5e-04 | 0.0432953 | 44.3624 | 15.4716 | -1.51972 | -2.08741 | KB882086.1:3066893-3074937 |
| XLOC_013843 | ENSAMXG00000021040 | 3.00057 | 9.78594 | 0.00015 | 0.0178305 | 3.00057 | 9.78594 | 1.70548 | 2.32338 | KB882086.1:2962328-2965271 |
| XLOC_013915 | cacna2d1a | 9.12871 | 3.60254 | 0.00045 | 0.0403574 | 9.12871 | 3.60254 | -1.3414 | -2.04831 | KB882087.1:3261405-3323962 |
| XLOC_013950 | cldn18 | 14.3133 | 87.1471 | 5e-05 | 0.00738566 | 14.3133 | 87.1471 | 2.6061 | 2.96862 | KB882088.1:2214646-2223396 |
| XLOC_014159 | ENSAMXG00000010068 | 43.151 | 16.2052 | 0.00075 | 0.0540027 | 43.151 | 16.2052 | -1.41294 | -1.93706 | KB882090.1:1681088-1682458 |
| XLOC_014216 | ENSAMXG00000010163 | 0 | 2.3057 | 5e-05 | 0.00738566 | 0 | 2.3057 | Inf | NA | KB882090.1:1920322-1924962 |
| XLOC_014269 | tdo2a | 9.30809 | 2.90932 | 0.00025 | 0.0254937 | 9.30809 | 2.90932 | -1.6778 | -2.17427 | KB882091.1:489980-497676 |

| Locus | gene_short_name | Dark(FPKM) | Light(FPKM) | Light_Dark_p_value | Light_Dark_q_value | Light_Dark_value_1 | Light_Dark_value_2 | Light_Dark_log2_fold_change | Light_Dark_test_stat | length |
|---|---|---|---|---|---|---|---|---|---|---|
| XLOC_014425 | fkbp9 | 25.2509 | 5.74246 | 5e-05 | 0.00738566 | 25.2509 | 5.74246 | -2.13659 | -3.32982 | KB882093.1:646064-654177 |
| XLOC_014455 | ENSAMXG00000028818 | 46.0104 | 6.83705 | 5e-05 | 0.00738566 | 46.0104 | 6.83705 | -2.75051 | -3.85869 | KB882094.1:71774-82002 |
| XLOC_014633 | grp | 1.91106 | 9.94569 | 5e-05 | 0.00738566 | 1.91106 | 9.94569 | 2.37969 | 2.92764 | KB882096.1:1162040-1168823 |
| XLOC_014802 | tmem82 | 5.41818 | 1.85308 | 0.00085 | 0.0574938 | 5.41818 | 1.85308 | -1.54788 | -2.01365 | KB882098.1:305735-310064 |
| XLOC_014878 | map1lc3cl | 6.03962 | 21.366 | 5e-05 | 0.00738566 | 6.03962 | 21.366 | 1.82278 | 2.54464 | KB882099.1:90771-93011 |
| XLOC_015020 | duox | 6.37988 | 36.0034 | 5e-05 | 0.00738566 | 6.37988 | 36.0034 | 2.49653 | 2.90409 | KB882100.1:2000619-2019230 |
| XLOC_015028 | asb15a | 3.41135 | 1.00845 | 0.00025 | 0.0254937 | 3.41135 | 1.00845 | -1.7582 | -2.3118 | KB882100.1:2710426-2716943 |
| XLOC_015067 | rab3da | 10.5606 | 23.8306 | 0.00135 | 0.0831906 | 10.5606 | 23.8306 | 1.17413 | 1.87603 | KB882101.1:1950042-1972519 |
| XLOC_015086 | ENSAMXG00000000481 | 7.68308 | 2.01204 | 3e-04 | 0.0296882 | 7.68308 | 2.01204 | -1.93302 | -2.26432 | KB882101.1:308357-310769 |
| XLOC_015239 | ENSAMXG00000005638 | 2.1106 | 0 | 5e-05 | 0.00738566 | 2.1106 | 0 | -Inf | NA | KB882103.1:688962-689632 |
| XLOC_015368 | rsph14 | 7.61391 | 1.25619 | 5e-05 | 0.00738566 | 7.61391 | 1.25619 | -2.59958 | -3.00133 | KB882105.1:515647-520846 |
| XLOC_015415 | si:busm1-266f07.2 | 55.2484 | 5.36906 | 5e-05 | 0.00738566 | 55.2484 | 5.36906 | -3.36319 | -4.58697 | KB882105.1:212135-221320 |
| XLOC_015421 | star | 1.22599 | 4.77582 | 5e-05 | 0.00738566 | 1.22599 | 4.77582 | 1.9618 | 2.448 | KB882105.1:400138-404684 |
| XLOC_015427 | si:ch211-170d8.5 | 3.81639 | 0.930637 | 2e-04 | 0.0216011 | 3.81639 | 0.930637 | -2.03592 | -2.4455 | KB882105.1:1559551-1568711 |
| XLOC_015638 | zgc:162144 | 3.89968 | 1.25653 | 8e-04 | 0.0556097 | 3.89968 | 1.25653 | -1.63392 | -2.06812 | KB882108.1:306368-308320 |
| XLOC_015734 | slc8a3 | 4.78051 | 1.85809 | 0.0014 | 0.0852261 | 4.78051 | 1.85809 | -1.36335 | -1.91299 | KB882109.1:2933085-3008501 |

| Locus | gene_short_name | Dark(FPKM) | Light(FPKM) | Light_Dark_p_value | Light_Dark_q_value | Light_Dark_value_1 | Light_Dark_value_2 | Light_Dark_log2_fold_change | Light_Dark_test_stat | length |
|---|---|---|---|---|---|---|---|---|---|---|
| XLOC_015751 | ENSAMXG00000011560 | 9.54339 | 42.8343 | 1e-04 | 0.0124776 | 9.54339 | 42.8343 | 2.16619 | 2.53697 | KB882110.1:206963-208029 |
| XLOC_015769 | ENSAMXG00000028550 | 24.6941 | 134.174 | 3e-04 | 0.0296882 | 24.6941 | 134.174 | 2.44187 | 4.5531 | KB882110.1:2696591-2697884 |
| XLOC_015813 | ENSAMXG00000007189 | 2114.1 | 7716.11 | 7e-04 | 0.0518904 | 2114.1 | 7716.11 | 1.86783 | 1.98399 | KB882111.1:1213159-1218786 |
| XLOC_016155 | calcoco1b | 0.168095 | 1.89092 | 5e-05 | 0.00738566 | 0.168095 | 1.89092 | 3.49174 | 3.55109 | KB882116.1:1649691-1665179 |
| XLOC_016157 | nr1d4b | 9.43547 | 3.81497 | 0.00065 | 0.0502225 | 9.43547 | 3.81497 | -1.30642 | -2.02862 | KB882116.1:1829292-1850489 |
| XLOC_016289 | nkx6.2 | 2.00559 | 9.50662 | 5e-05 | 0.00738566 | 2.00559 | 9.50662 | 2.2449 | 2.95916 | KB882118.1:27043-29344 |
| XLOC_016404 | cabp5a | 11.0573 | 25.4383 | 0.00115 | 0.0742841 | 11.0573 | 25.4383 | 1.202 | 1.89508 | KB882119.1:203599-209419 |
| XLOC_016478 | nox1 | 7.13281 | 21.4823 | 5e-05 | 0.00738566 | 7.13281 | 21.4823 | 1.5906 | 2.3422 | KB882119.1:2377781-2392325 |
| XLOC_016636 | zgc:112408 | 3.10042 | 14.8977 | 1e-04 | 0.0124776 | 3.10042 | 14.8977 | 2.26455 | 2.70322 | KB882121.1:1709196-1714770 |
| XLOC_016812 | ENSAMXG00000025425 | 6.69814 | 33.1008 | 0.00125 | 0.0789662 | 6.69814 | 33.1008 | 2.30503 | 2.46293 | KB882124.1:1942746-1943496 |
| XLOC_016815 | rida | 34.3824 | 13.6325 | 4e-04 | 0.0372019 | 34.3824 | 13.6325 | -1.33462 | -2.034 | KB882124.1:2108955-2115104 |
| XLOC_016913 | prom2 | 11.7115 | 26.8014 | 0.00145 | 0.0861806 | 11.7115 | 26.8014 | 1.19438 | 1.90546 | KB882125.1:851909-874460 |
| XLOC_016970 | ENSAMXG00000012464 | 5.5153 | 1.34483 | 5e-05 | 0.00738566 | 5.5153 | 1.34483 | -2.03601 | -2.62552 | KB882125.1:2097923-2107121 |
| XLOC_016987 | si:dkey-61p9.11 | 4.52351 | 1.33199 | 1e-04 | 0.0124776 | 4.52351 | 1.33199 | -1.76386 | -2.39382 | KB882126.1:318388-326998 |
| XLOC_017269 | rtn2b | 21.6673 | 8.93054 | 0.00065 | 0.0502225 | 21.6673 | 8.93054 | -1.2787 | -1.98303 | KB882129.1:352176-365346 |
| XLOC_017677 | CRACR2A | 1.12663 | 3.77506 | 0.00055 | 0.0450978 | 1.12663 | 3.77506 | 1.74449 | 2.28683 | KB882135.1:97375-125255 |

| Locus | gene_short_name | Dark(FPKM) | Light(FPKM) | Light_Dark_p_value | Light_Dark_q_value | Light_Dark_value_1 | Light_Dark_value_2 | Light_Dark_log2_fold_change | Light_Dark_test_stat | length |
|---|---|---|---|---|---|---|---|---|---|---|
| XLOC_017764 | slc1a7a | 2.09096 | 0.527558 | 2e-04 | 0.0216011 | 2.09096 | 0.527558 | -1.98676 | -2.39707 | KB882137.1:887543-921811 |
| XLOC_017952 | ENSAMXG00000018683 | 7.53352 | 2.57226 | 2e-04 | 0.0216011 | 7.53352 | 2.57226 | -1.55029 | -2.2039 | KB882139.1:2130673-2153246 |
| XLOC_017953 | ceacam1 | 8.10879 | 42.4682 | 5e-05 | 0.00738566 | 8.10879 | 42.4682 | 2.38883 | 2.99767 | KB882139.1:2170775-2187055 |
| XLOC_017991 | cnfn | 58.4859 | 236.735 | 0.00015 | 0.0178305 | 58.4859 | 236.735 | 2.01711 | 2.33068 | KB882139.1:2229954-2231596 |
| XLOC_018093 | rcn3 | 13.974 | 3.02097 | 5e-05 | 0.00738566 | 13.974 | 3.02097 | -2.20966 | -3.20665 | KB882142.1:1293779-1298434 |
| XLOC_018150 | ENSAMXG00000010386 | 1.1248 | 0 | 5e-05 | 0.00738566 | 1.1248 | 0 | -Inf | NA | KB882142.1:1634996-1656339 |
| XLOC_018187 | noxo1b | 6.97124 | 70.0027 | 5e-05 | 0.00738566 | 6.97124 | 70.0027 | 3.32792 | 4.0743 | KB882143.1:1198526-1206548 |
| XLOC_018209 | tefa | 14.5185 | 32.6502 | 7e-04 | 0.0518904 | 14.5185 | 32.6502 | 1.1692 | 1.91382 | KB882143.1:2307401-2311742 |
| XLOC_018289 | lrrc58a | 1.30221 | 3.77552 | 0.00135 | 0.0831906 | 1.30221 | 3.77552 | 1.53571 | 1.89232 | KB882144.1:970506-980419 |
| XLOC_018397 | p3h1 | 4.94728 | 2.12861 | 0.001 | 0.0665199 | 4.94728 | 2.12861 | -1.21672 | -1.84707 | KB882145.1:2066286-2082635 |
| XLOC_018467 | esrrd | 4.05731 | 12.2426 | 5e-05 | 0.00738566 | 4.05731 | 12.2426 | 1.59332 | 2.46542 | KB882146.1:1513311-1526512 |
| XLOC_018486 | ENSAMXG00000020645 | 393.498 | 141.315 | 5e-04 | 0.0432953 | 393.498 | 141.315 | -1.47744 | -2.18495 | KB882146.1:1286623-1287664 |
| XLOC_018494 | foxa3 | 6.76531 | 15.3178 | 0.0013 | 0.0811047 | 6.76531 | 15.3178 | 1.17898 | 1.87088 | KB882146.1:1859363-1865147 |
| XLOC_018523 | rgcc | 33.705 | 98.6289 | 5e-05 | 0.00738566 | 33.705 | 98.6289 | 1.54905 | 2.50061 | KB882147.1:1553242-1559475 |
| XLOC_018615 | ENSAMXG00000020762 | 2.54283 | 0.711618 | 0.00145 | 0.0861806 | 2.54283 | 0.711618 | -1.83726 | -2.05134 | KB882149.1:354205-419865 |
| XLOC_018701 | clic5a | 4.24356 | 1.59612 | 7e-04 | 0.0518904 | 4.24356 | 1.59612 | -1.4107 | -1.9045 | KB882150.1:1879199-1890582 |

| Locus | gene_short_name | Dark(FPKM) | Light(FPKM) | Light_Dark_p_value | Light_Dark_q_value | Light_Dark_value_1 | Light_Dark_value_2 | Light_Dark_log2_fold_change | Light_Dark_test_stat | length |
|---|---|---|---|---|---|---|---|---|---|---|
| XLOC_018792 | ENSAMXG00000009725 | 18.941 | 51.8438 | 3e-04 | 0.0296882 | 18.941 | 51.8438 | 1.45266 | 2.07903 | KB882151.1:2828307-2836759 |
| XLOC_018793 | APOH | 8.92664 | 3.40406 | 0.0015 | 0.0878528 | 8.92664 | 3.40406 | -1.39086 | -1.83549 | KB882152.1:451566-468120 |
| XLOC_018827 | aanat1 | 27.3703 | 4.58755 | 5e-05 | 0.00738566 | 27.3703 | 4.58755 | -2.57682 | -3.37155 | KB882152.1:1429795-1431775 |
| XLOC_018850 | rsad2 | 2.33004 | 10.4055 | 5e-05 | 0.00738566 | 2.33004 | 10.4055 | 2.15892 | 2.77737 | KB882153.1:1732446-1736226 |
| XLOC_019129 | fetub | 19.1377 | 6.40619 | 1e-04 | 0.0124776 | 19.1377 | 6.40619 | -1.57888 | -2.23955 | KB882159.1:316425-328721 |
| XLOC_019196 | tm6sf2 | 8.66805 | 3.00263 | 1e-04 | 0.0124776 | 8.66805 | 3.00263 | -1.52948 | -2.34197 | KB882160.1:2065541-2087059 |
| XLOC_019233 | slc5a8 | 0.618203 | 2.59508 | 0.00065 | 0.0502225 | 0.618203 | 2.59508 | 2.06962 | 2.11096 | KB882162.1:1869266-1878015 |
| XLOC_019321 | tnni4b.2 | 72.4225 | 28.0282 | 0.00065 | 0.0502225 | 72.4225 | 28.0282 | -1.36956 | -2.06959 | KB882164.1:69935-72299 |
| XLOC_019391 | ENSAMXG00000009469 | 305.971 | 89.4519 | 7e-04 | 0.0518904 | 305.971 | 89.4519 | -1.77421 | -2.1061 | KB882165.1:1308251-1310558 |
| XLOC_019393 | ENSAMXG00000009481 | 24.4729 | 6.44703 | 0.00065 | 0.0502225 | 24.4729 | 6.44703 | -1.92448 | -2.03948 | KB882165.1:1330064-1348296 |
| XLOC_019441 | me1 | 68.1913 | 18.7993 | 5e-05 | 0.00738566 | 68.1913 | 18.7993 | -1.85891 | -2.75406 | KB882166.1:2099378-2196189 |
| XLOC_019461 | ENSAMXG00000008750 | 2.75925 | 14.4991 | 5e-05 | 0.00738566 | 2.75925 | 14.4991 | 2.39361 | 3.82746 | KB882166.1:1824805-1848177 |
| XLOC_019467 | ENSAMXG00000008922 | 1.64697 | 5.64861 | 3e-04 | 0.0296882 | 1.64697 | 5.64861 | 1.77809 | 2.41595 | KB882166.1:2531268-2533496 |
| XLOC_019560 | si:ch211-63p21.8 | 0.384448 | 1.54061 | 0.00015 | 0.0178305 | 0.384448 | 1.54061 | 2.00264 | 2.54532 | KB882169.1:123635-128754 |
| XLOC_019685 | igfn1.3 | 5.57065 | 2.14095 | 0.00015 | 0.0178305 | 5.57065 | 2.14095 | -1.37959 | -2.13406 | KB882171.1:4941586-4966155 |
| XLOC_019690 | ptgis | 0.797064 | 2.39421 | 0.00075 | 0.0540027 | 0.797064 | 2.39421 | 1.58678 | 1.95254 | KB882171.1:5206751-5226126 |

| Locus | gene_short_name | Dark(FPKM) | Light(FPKM) | Light_Dark_p_value | Light_Dark_q_value | Light_Dark_value_1 | Light_Dark_value_2 | Light_Dark_log2_fold_change | Light_Dark_test_stat | length |
|---|---|---|---|---|---|---|---|---|---|---|
| XLOC_019772 | rbp7b | 37.7257 | 9.62923 | 5e-05 | 0.00738566 | 37.7257 | 9.62923 | -1.97005 | -3.11842 | KB882171.1:5730311-5734124 |
| XLOC_019868 | vill | 5.25876 | 22.0608 | 5e-05 | 0.00738566 | 5.25876 | 22.0608 | 2.06869 | 2.66002 | KB882174.1:2304249-2332392 |
| XLOC_019943 | CLDN18 | 26.2224 | 183.058 | 1e-04 | 0.0124776 | 26.2224 | 183.058 | 2.80343 | 2.94561 | KB882176.1:135549-140398 |
| XLOC_019971 | ENSAMXG00000005446 | 8.47209 | 3.49831 | 8e-04 | 0.0556097 | 8.47209 | 3.49831 | -1.27606 | -1.95463 | KB882177.1:430529-441690 |
| XLOC_019973 | ENSAMXG00000028341 | 3.07469 | 1.03083 | 0.0015 | 0.0878528 | 3.07469 | 1.03083 | -1.57663 | -1.90166 | KB882177.1:508417-518105 |
| XLOC_019983 | frem3 | 6.19939 | 2.87584 | 0.00125 | 0.0789662 | 6.19939 | 2.87584 | -1.10814 | -1.81975 | KB882177.1:2087756-2162568 |
| XLOC_020107 | capn3b | 4.18702 | 1.09168 | 5e-05 | 0.00738566 | 4.18702 | 1.09168 | -1.93938 | -2.69238 | KB882181.1:493429-517067 |
| XLOC_020174 | zmp:0000000846 | 5.76813 | 28.0792 | 5e-05 | 0.00738566 | 5.76813 | 28.0792 | 2.28333 | 3.10672 | KB882182.1:289544-377076 |
| XLOC_020324 | si:dkey-183i3.5 | 116.963 | 44.9713 | 5e-04 | 0.0432953 | 116.963 | 44.9713 | -1.37898 | -1.98152 | KB882184.1:1833668-1842014 |
| XLOC_020338 | ghrl | 8.32656 | 46.2696 | 5e-05 | 0.00738566 | 8.32656 | 46.2696 | 2.47427 | 3.16788 | KB882185.1:955034-956773 |
| XLOC_020358 | fkbp5 | 3.16764 | 12.8911 | 5e-05 | 0.00738566 | 3.16764 | 12.8911 | 2.02489 | 2.87151 | KB882185.1:1474306-1492367 |
| XLOC_020421 | f9a | 1.79665 | 0.510339 | 5e-04 | 0.0432953 | 1.79665 | 0.510339 | -1.81578 | -2.18805 | KB882186.1:298362-306971 |
| XLOC_020515 | trhra | 0.557703 | 1.51277 | 0.0016 | 0.0923632 | 0.557703 | 1.51277 | 1.43963 | 1.88077 | KB882188.1:127702-138105 |
| XLOC_020582 | zgc:172270 | 10.5903 | 4.37526 | 0.0015 | 0.0878528 | 10.5903 | 4.37526 | -1.27531 | -1.81519 | KB882189.1:407700-411259 |
| XLOC_020691 | RF00411 | 0 | 36.4129 | 0.00145 | 0.0861806 | 0 | 36.4129 | Inf | NA | KB882192.1:1879264-1879400 |
| XLOC_020745 | ptgdsb.1 | 46.8369 | 186.845 | 5e-05 | 0.00738566 | 46.8369 | 186.845 | 1.99612 | 3.19167 | KB882193.1:1400216-1402995 |

| Locus | gene_short_name | Dark(FPKM) | Light(FPKM) | Light_Dark_p_value | Light_Dark_q_value | Light_Dark_value_1 | Light_Dark_value_2 | Light_Dark_log2_fold_change | Light_Dark_test_stat | length |
|---|---|---|---|---|---|---|---|---|---|---|
| XLOC_020783 | f10 | 10.4939 | 3.58352 | 0.00045 | 0.0403574 | 10.4939 | 3.58352 | -1.5501 | -2.12136 | KB882194.1:1862976-1873343 |
| XLOC_020842 | ENSAMXG00000020478 | 1.33991 | 5.45259 | 4e-04 | 0.0372019 | 1.33991 | 5.45259 | 2.02481 | 2.29157 | KB882196.1:160868-173864 |
| XLOC_021022 | myoz2a | 32.0886 | 9.48206 | 5e-05 | 0.00738566 | 32.0886 | 9.48206 | -1.75879 | -2.52943 | KB882199.1:158356-174366 |
| XLOC_021193 | ENSAMXG00000028390 | 25.5123 | 8.99533 | 0.00055 | 0.0450978 | 25.5123 | 8.99533 | -1.50395 | -2.01482 | KB882202.1:2073617-2084028 |
| XLOC_021216 | si:zfos-223e1.2 | 6.32949 | 0.938575 | 5e-05 | 0.00738566 | 6.32949 | 0.938575 | -2.75355 | -3.89346 | KB882202.1:3836973-3885323 |
| XLOC_021299 | vtcn1 | 1.57467 | 12.2831 | 5e-05 | 0.00738566 | 1.57467 | 12.2831 | 2.96355 | 3.30079 | KB882203.1:2048678-2052087 |
| XLOC_021346 | EIF5AL1 | 64.8907 | 27.7358 | 0.0014 | 0.0852261 | 64.8907 | 27.7358 | -1.22626 | -1.90473 | KB882205.1:761165-766595 |
| XLOC_021415 | hpda | 33.1709 | 10.1052 | 2e-04 | 0.0216011 | 33.1709 | 10.1052 | -1.71482 | -2.3654 | KB882207.1:1071964-1110817 |
| XLOC_021445 | hspb1 | 58.7202 | 25.5665 | 7e-04 | 0.0518904 | 58.7202 | 25.5665 | -1.1996 | -1.93777 | KB882207.1:1632877-1642891 |
| XLOC_021617 | agt | 5.39916 | 1.87453 | 0.00085 | 0.0574938 | 5.39916 | 1.87453 | -1.52621 | -2.00331 | KB882210.1:608040-616510 |
| XLOC_021679 | gls2a | 6.02413 | 13.7149 | 0.0011 | 0.0719801 | 6.02413 | 13.7149 | 1.18692 | 1.83241 | KB882211.1:1237783-1270711 |
| XLOC_021786 | slc16a7 | 2.23961 | 0.429028 | 0.00055 | 0.0450978 | 2.23961 | 0.429028 | -2.3841 | -2.49175 | KB882214.1:536854-545219 |
| XLOC_021793 | si:ch73-269m23.5 | 5.17944 | 1.71751 | 0.00055 | 0.0450978 | 5.17944 | 1.71751 | -1.59248 | -2.13826 | KB882214.1:9545-14471 |
| XLOC_021804 | krt5 | 557.563 | 196.279 | 0.00075 | 0.0540027 | 557.563 | 196.279 | -1.50623 | -1.92066 | KB882214.1:724231-728166 |
| XLOC_021915 | dbpb | 4.68001 | 0.931464 | 5e-05 | 0.00738566 | 4.68001 | 0.931464 | -2.32894 | -3.0408 | KB882219.1:812678-823637 |
| XLOC_021990 | tdh | 28.7533 | 74.4462 | 0.00075 | 0.0540027 | 28.7533 | 74.4462 | 1.37247 | 2.07401 | KB882221.1:418728-427626 |

| Locus | gene_short_name | Dark(FPKM) | Light(FPKM) | Light_Dark_p_value | Light_Dark_q_value | Light_Dark_value_1 | Light_Dark_value_2 | Light_Dark_log2_fold_change | Light_Dark_test_stat | length |
|---|---|---|---|---|---|---|---|---|---|---|
| XLOC_022058 | cry1bb | 7.1691 | 16.9325 | 4e-04 | 0.0372019 | 7.1691 | 16.9325 | 1.23993 | 2.01126 | KB882222.1:1131911-1144602 |
| XLOC_022096 | apobec2a | 74.6589 | 25.1353 | 5e-05 | 0.00738566 | 74.6589 | 25.1353 | -1.5706 | -2.45778 | KB882222.1:927578-928665 |
| XLOC_022122 | kng1 | 33.3538 | 12.3665 | 0.00105 | 0.0689328 | 33.3538 | 12.3665 | -1.43141 | -1.84292 | KB882223.1:700652-707023 |
| XLOC_022134 | ENSAMXG00000001567 | 12.9239 | 3.85645 | 5e-05 | 0.00738566 | 12.9239 | 3.85645 | -1.74469 | -2.42787 | KB882223.1:693259-698500 |
| XLOC_022209 | ENSAMXG00000011255 | 3.46259 | 0.791043 | 1e-04 | 0.0124776 | 3.46259 | 0.791043 | -2.13002 | -2.38538 | KB882226.1:586876-592871 |
| XLOC_022287 | ENSAMXG00000012686 | 21.9871 | 5.11093 | 5e-05 | 0.00738566 | 21.9871 | 5.11093 | -2.105 | -2.83134 | KB882227.1:978239-982001 |
| XLOC_022304 | ttr | 23.5395 | 9.2831 | 0.00055 | 0.0450978 | 23.5395 | 9.2831 | -1.3424 | -1.91798 | KB882228.1:265590-269296 |
| XLOC_022388 | nfil3-5 | 12.51 | 42.5922 | 5e-05 | 0.00738566 | 12.51 | 42.5922 | 1.76751 | 2.88125 | KB882230.1:1520805-1522581 |
| XLOC_022467 | dhrs7cb | 36.8643 | 12.793 | 5e-05 | 0.00738566 | 36.8643 | 12.793 | -1.52687 | -2.43824 | KB882231.1:1041292-1055122 |
| XLOC_022521 | ENSAMXG00000014315 | 7.85082 | 1.96306 | 5e-05 | 0.00738566 | 7.85082 | 1.96306 | -1.99974 | -2.65794 | KB882233.1:3476411-3489654 |
| XLOC_022604 | PLCB | 1.16374 | 3.26331 | 5e-05 | 0.00738566 | 1.16374 | 3.26331 | 1.48757 | 2.18254 | KB882233.1:6157952-6212655 |
| XLOC_022632 | cbln17 | 25.974 | 6.97346 | 0.00065 | 0.0502225 | 25.974 | 6.97346 | -1.89712 | -2.14317 | KB882234.1:1382594-1388624 |
| XLOC_022748 | IMPG2 | 4.8247 | 1.50829 | 1e-04 | 0.0124776 | 4.8247 | 1.50829 | -1.67753 | -2.36292 | KB882237.1:1046210-1099373 |
| XLOC_022789 | anxa6 | 59.7876 | 25.8033 | 8e-04 | 0.0556097 | 59.7876 | 25.8033 | -1.21229 | -1.86622 | KB882239.1:846264-881430 |
| XLOC_022881 | tfa | 211.861 | 60.2335 | 0.00045 | 0.0403574 | 211.861 | 60.2335 | -1.81448 | -2.16776 | KB882241.1:1709775-1723302 |
| XLOC_023022 | casq1a | 25.6961 | 7.42792 | 5e-05 | 0.00738566 | 25.6961 | 7.42792 | -1.79052 | -2.63776 | KB882246.1:808530-816311 |

| Locus | gene_short_name | Dark(FPKM) | Light(FPKM) | Light_Dark_p_value | Light_Dark_q_value | Light_Dark_value_1 | Light_Dark_value_2 | Light_Dark_log2_fold_change | Light_Dark_test_stat | length |
|---|---|---|---|---|---|---|---|---|---|---|
| XLOC_023119 | fah | 26.7036 | 12.4046 | 0.00165 | 0.0947053 | 26.7036 | 12.4046 | -1.10616 | -1.81082 | KB882249.1:1683760-1717112 |
| XLOC_023153 | asb2a.1 | 0.513023 | 2.7796 | 0.00025 | 0.0254937 | 0.513023 | 2.7796 | 2.43778 | 2.67106 | KB882250.1:1110546-1115791 |
| XLOC_023194 | si:ch211-232m10.6 | 0.842842 | 2.38 | 0.00065 | 0.0502225 | 0.842842 | 2.38 | 1.49763 | 2.01653 | KB882251.1:861514-871137 |
| XLOC_023206 | ENSAMXG00000009598 | 234.387 | 59.699 | 5e-05 | 0.00738566 | 234.387 | 59.699 | -1.97311 | -2.71601 | KB882251.1:1492607-1500271 |
| XLOC_023301 | ARL14 | 4.83578 | 26.1987 | 0.00065 | 0.0502225 | 4.83578 | 26.1987 | 2.43767 | 2.48397 | KB882254.1:1191647-1192262 |
| XLOC_023352 | trim63a | 18.3331 | 2.05562 | 5e-05 | 0.00738566 | 18.3331 | 2.05562 | -3.1568 | -4.49833 | KB882256.1:514228-515290 |
| XLOC_023415 | ENSAMXG00000028514 | 1.80512 | 0 | 0.00035 | 0.0338036 | 1.80512 | 0 | -Inf | NA | KB882257.1:897719-899753 |
| XLOC_023576 | camk1gb | 7.04359 | 21.5279 | 1e-04 | 0.0124776 | 7.04359 | 21.5279 | 1.61183 | 2.49612 | KB882262.1:585977-605973 |
| XLOC_023658 | ENSAMXG00000025008 | 11.4043 | 41.6757 | 5e-05 | 0.00738566 | 11.4043 | 41.6757 | 1.86962 | 2.5223 | KB882265.1:1479264-1479678 |
| XLOC_023659 | ENSAMXG00000020401 | 13.8796 | 1.62433 | 0.00055 | 0.0450978 | 13.8796 | 1.62433 | -3.09505 | -3.14221 | KB882265.1:1502164-1502865 |
| XLOC_023687 | fbxo21 | 5.25162 | 14.739 | 5e-05 | 0.00738566 | 5.25162 | 14.739 | 1.4888 | 2.29414 | KB882266.1:1730495-1770191 |
| XLOC_023762 | si:ch1073-440b2.1 | 10.4369 | 25.4015 | 0.00045 | 0.0403574 | 10.4369 | 25.4015 | 1.28322 | 2.07947 | KB882269.1:246386-274675 |
| XLOC_023791 | si:dkey-220f10.4 | 3.82907 | 1.2969 | 0.00055 | 0.0450978 | 3.82907 | 1.2969 | -1.56192 | -2.05953 | KB882270.1:302818-310003 |
| XLOC_023852 | zgc:158404 | 13.399 | 4.48923 | 5e-05 | 0.00738566 | 13.399 | 4.48923 | -1.57758 | -2.36692 | KB882270.1:4030758-4041129 |
| XLOC_023950 | enpp7.2 | 5.34996 | 16.4123 | 5e-05 | 0.00738566 | 5.34996 | 16.4123 | 1.61718 | 2.38606 | KB882270.1:5657277-5668243 |
| XLOC_023962 | rs1a | 17.4585 | 43.4856 | 0.00015 | 0.0178305 | 17.4585 | 43.4856 | 1.31661 | 2.12548 | KB882271.1:45462-56630 |

| Locus | gene_short_name | Dark(FPKM) | Light(FPKM) | Light_Dark_p_value | Light_Dark_q_value | Light_Dark_value_1 | Light_Dark_value_2 | Light_Dark_log2_fold_change | Light_Dark_test_stat | length |
|---|---|---|---|---|---|---|---|---|---|---|
| XLOC_023975 | si:dkey-18a10.3 | 7.40938 | 16.9341 | 0.00095 | 0.0638279 | 7.40938 | 16.9341 | 1.19251 | 1.84885 | KB882272.1:193251-203983 |
| XLOC_024005 | myl10 | 331.775 | 112.53 | 0.00135 | 0.0831906 | 331.775 | 112.53 | -1.5599 | -2.01332 | KB882272.1:489176-491899 |
| XLOC_024058 | si:dkey-183n20.15 | 9.33295 | 3.21923 | 1e-04 | 0.0124776 | 9.33295 | 3.21923 | -1.53562 | -2.03294 | KB882273.1:978216-986042 |
| XLOC_024060 | cyp2ad6 | 2.53474 | 7.96732 | 2e-04 | 0.0216011 | 2.53474 | 7.96732 | 1.65226 | 2.20046 | KB882273.1:1043530-1050357 |
| XLOC_024365 | abcb11a | 5.76304 | 0.680922 | 5e-05 | 0.00738566 | 5.76304 | 0.680922 | -3.08127 | -4.13732 | KB882284.1:888672-900259 |
| XLOC_024373 | ENSAMXG00000006171 | 10.6621 | 25.7904 | 8e-04 | 0.0556097 | 10.6621 | 25.7904 | 1.27435 | 1.95357 | KB882284.1:1287431-1292320 |
| XLOC_024375 | ENSAMXG00000005569 | 9.14128 | 34.7095 | 5e-05 | 0.00738566 | 9.14128 | 34.7095 | 1.92486 | 2.94981 | KB882284.1:390628-401042 |
| XLOC_024379 | TUBA4A | 99.7674 | 15.5236 | 5e-05 | 0.00738566 | 99.7674 | 15.5236 | -2.6841 | -4.09711 | KB882284.1:504439-507144 |
| XLOC_024386 | klhl41a | 8.44525 | 2.87167 | 0.00065 | 0.0502225 | 8.44525 | 2.87167 | -1.55625 | -2.1465 | KB882284.1:741078-746501 |
| XLOC_024388 | col28a2a | 2.70993 | 0.611707 | 5e-05 | 0.00738566 | 2.70993 | 0.611707 | -2.14735 | -3.05323 | KB882284.1:754399-776147 |
| XLOC_024488 | trpa1b | 0.608163 | 2.41025 | 1e-04 | 0.0124776 | 0.608163 | 2.41025 | 1.98665 | 2.5676 | KB882288.1:781656-839392 |
| XLOC_024551 | rgs9bp | 1.09637 | 3.81565 | 5e-05 | 0.00738566 | 1.09637 | 3.81565 | 1.79919 | 2.49079 | KB882290.1:1513855-1522129 |
| XLOC_024582 | UBAP1L | 3.72568 | 0.708624 | 5e-05 | 0.00738566 | 3.72568 | 0.708624 | -2.39441 | -3.11869 | KB882292.1:694761-714495 |
| XLOC_024590 | ENSAMXG00000003434 | 18.931 | 52.7914 | 8e-04 | 0.0556097 | 18.931 | 52.7914 | 1.47956 | 2.04781 | KB882292.1:1414437-1474066 |
| XLOC_024818 | VSIG2 | 2.34237 | 12.6587 | 7e-04 | 0.0518904 | 2.34237 | 12.6587 | 2.43409 | 2.46938 | KB882299.1:1085728-1098687 |
| XLOC_025145 | rnasel3 | 73.7627 | 22.5763 | 1e-04 | 0.0124776 | 73.7627 | 22.5763 | -1.70808 | -2.54725 | KB882311.1:1429665-1430118 |

| Locus | gene_short_name | Dark(FPKM) | Light(FPKM) | Light_Dark_p_value | Light_Dark_q_value | Light_Dark_value_1 | Light_Dark_value_2 | Light_Dark_log2_fold_change | Light_Dark_test_stat | length |
|---|---|---|---|---|---|---|---|---|---|---|
| XLOC_025152 | ENSAMXG00000001701 | 145.931 | 51.2912 | 5e-05 | 0.00738566 | 145.931 | 51.2912 | -1.5085 | -2.33554 | KB882311.1:471525-477166 |
| XLOC_025153 | ENSAMXG00000001722 | 29.1152 | 217.014 | 5e-05 | 0.00738566 | 29.1152 | 217.014 | 2.89795 | 4.3711 | KB882311.1:481635-486402 |
| XLOC_025154 | apoa1a | 208.682 | 54.5999 | 5e-05 | 0.00738566 | 208.682 | 54.5999 | -1.93434 | -2.89141 | KB882311.1:585779-587980 |
| XLOC_025169 | SCTR | 1.89309 | 8.1624 | 5e-05 | 0.00738566 | 1.89309 | 8.1624 | 2.10825 | 3.05091 | KB882312.1:152821-170018 |

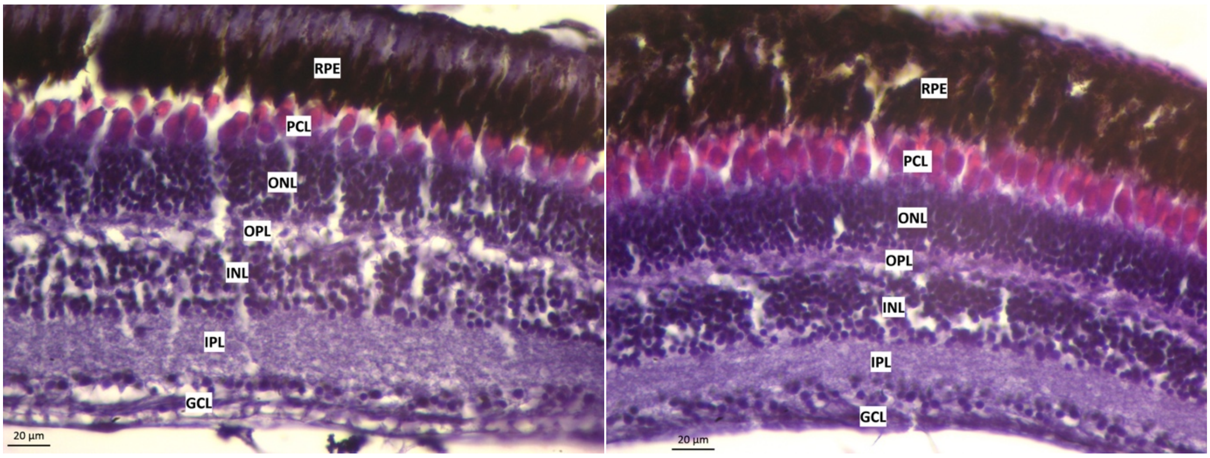


Supplementary Figure 1. Cross section of retinal layers in surface fish reared in L/D (top) or D/D (bottom) conditions. GCL - ganglion cell layer; IPL - inner plexiform layer; INL - inner nuclear layer; OPL - outer plexiform layer; ONL - outer nuclear layer; PCL - photoreceptor cell layer; RPE - retinal pigment epithelium. Scale bar 20 µm.

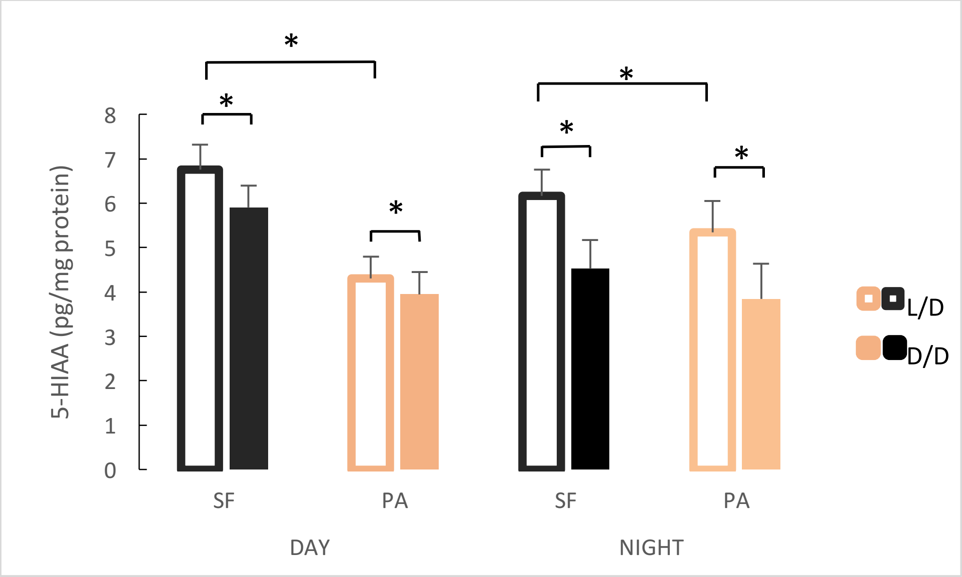


Supplementary Figure 2. Levels of 5-Hydroxyindoleacetic acid (5-HIAA), the main metabolite of serotonin, in adult brains of L/D or D/D-reared surface fish (SF) and Pachón cavefish (PA) collected in the middle of the day (DAY) and the middle of the night (NIGHT). (Error bars represent the standard error of the means.)
